# Identifying a novel mechanism of L-leucine uptake in *Mycobacterium tuberculosis* using a chemical genomic approach

**DOI:** 10.1101/2025.04.14.648691

**Authors:** Nisheeth Agarwal, Himanshu Gogoi, Eeba, Linus Augustin, Md. Younus Khan, Yashwant Kumar, Sayan Kumar Bhowmick, Bappaditya Dey

## Abstract

Amino acid biosynthesis is vital for *Mycobacterium tuberculosis* (Mtb) proliferation and tuberculosis pathogenesis. However, it is not clear how amino acids are transported in Mtb, particularly the branched chain amino acids (BCAAs) that contribute to the production of the cell-wall lipid component precursors such as acetyl-CoA and propionyl-CoA.

While performing the screening of an FDA-approved repurposed library of small molecule inhibitors against the auxotrophic strain Mtb mc^2^ 6206, which lacks *leuC-leuD* and *panC-panD* genes, we identified a molecule namely semapimod, which exclusively inhibits growth of the auxotrophic strain, whereas no effect is observed against the wild-type Mtb H_37_Rv. Interestingly, 24 h of exposure of Mtb mc^2^ 6206 to semapimod causes massive transcriptional reprogramming with differential expression of >450 genes associated with a myriad of metabolic activities. By performing a series of experiments, we affirm that semapimod indeed inhibits the L-leucine uptake in Mtb mc^2^ 6206 by targeting a protein involved in the cell-wall lipid biosynthesis pathway. Remarkably, semapimod treatment of mice infected with Mtb H_37_Rv causes a significant reduction of bacterial load in lungs and spleen, despite showing no efficacy against the pathogenic strain *in vitro*.

Overall findings of our study reveal that together with an endogenous pathway for L-leucine biosynthesis, a well-orchestrated machinery for its uptake is functional in Mtb which is important for intracellular survival of the TB pathogen.

## Introduction

Tuberculosis (TB), caused by the pathogen *Mycobacterium tuberculosis* (Mtb) is the leading cause of death due to microbial infections. As per the recent estimates, approximately 25% of the global population is asymptomatically infected with Mtb and about 11 million people developed the disease annually resulting in the loss of ∼1.25 million lives. In 2023, eight countries accounted for more than two thirds of the global total burden, with India leading the count^1^.

TB is curable and the usual treatment of pulmonary TB (PTB) cases involves 4 antibiotics (isoniazid (H), rifampicin(R), pyrazinamide (Z) and ethambutol (E)) for the first 2 months followed by four months of treatment with R and H^1, 2, 3, 4^. Unfortunately, due to multiple reasons such as delayed or mis-diagnosis, inadequate availability of drugs in resource-limited settings, poor administration, and noncompliance to the prolonged toxic drug regimen, TB becomes resistant to these drugs^5, 6, 7^. The drug-resistant TB (DR-TB) is classified under five categories: H-resistant TB^8^, R-resistant TB (RR-TB)^9^, multidrug-resistant TB (MDR-TB) which is resistant to both H and R, pre-extensively drug-resistant TB (pre-XDR-TB) which is resistant to H, R and any fluoroquinolone or any of the second-line injectables such as amikacin, capreomycin, and kanamycin, and XDR-TB which is resistant to H, R, any fluoroquinolone, and a second-line injectable or at least one of the recently developed drugs bedaquiline and linezolid. Treatment of DR-TB involves lengthy treatment with a course of second-line drugs for 6-18 months, causing huge socioeconomic burden^10^. DR-TB continues to be a threat to the public health system as number of people developing MDR/RR-TB remains stable since 2020. Globally, ∼400,000 MDR/RR-TB cases were reported in 2023, with about one fourth of the global MDR-TB cases reported from India alone^1, 11^. Considering the grim situation of everlasting DR-TB cases and emergence of resistance to newly developed drugs, there is a pressing need to explore novel therapeutic approaches.

With advancements in the medicinal chemistry, large numbers of new chemical entities (NCEs) have been synthesized over the last three decades that were evaluated for antimicrobial activity by the target-based and the phenotypic screening approaches^12, 13^. However, only a few of the newly developed antibiotics that got approval for human use exhibit a novel mechanism of action (MoA), whereas majority are the derivatives of the existing antibiotic classes^12, 14^. Evaluation of the antibiotic potential of the existing drugs that are approved for human use by the Federal Drug Agency (FDA) for other diseases, also known as drug repurposing, has emerged as an alternative approach to expedite treatment of infectious diseases including TB, as it avoids much of the hurdles at the early phases of drug development^12, 15^.

Herein, we report that a new anti-inflammatory molecule– semapimod, identified from a screen of the repurposed library of FDA-approved drugs, kills Mtb mc^2^ 6206 (a leucine and pantothenate auxotroph) at a reasonably low concentration of 20nM. To our great surprise, semapimod fails to inhibit other mycobacterial species including pathogenic Mtb H_37_Rv. These unexpected findings led us to identify a functional L-leucine uptake system in Mtb, which is blocked by the newly-identified compound. We show that semapimod targets an enzyme, PpsB, involved in the synthesis of the cell-wall phthiocerol dimycocerosate (PDIM), and influences Mtb virulence during host infection.

## Results

### Screening of FDA-approved library of molecules against Mtb mc^2^ 6206

In this study we screened a repurposed library (MedChemExpress) consisting of 3614 FDA-approved small molecule inhibitors against Mtb mc^2^ 6206, which is commonly used as a surrogate strain for pathogenic Mtb H_37_Rv^16^. These molecules primarily target microbial infections (790), GPCR/G Protein (646), apoptosis (371), metabolic enzyme (including protease) (355), membrane transporter (236), autophagy (210), cell cycle/DNA damage (157), immunology/inflammation (109), protein tyrosine kinase/RTK (99) and a large number of molecules targeting other miscellaneous pathways (402) (Supplementary Fig. 1a). The library contains a versatile set of molecules targeting different diseases such as cancer, cardiovascular disease, endocrinology, infection, inflammation/immunology, metabolic disease and neurological disease (Supplementary Fig. 1b). While these are intended for human use, 66% of the molecules have recently been launched, whereas remaining inhibitors are at different phases of trial, as depicted in Supplementary Fig. 1c. Of these, we identified 422 molecules showing complete inhibition of growth at 50µM concentration. As anticipated, majority (158) of the Mtb-inhibitors show anti-infection activity. However, several other compounds acting on host metabolic pathways, such as GPCR/G protein (68), apoptosis (45), autophagy (24), cell cycle/DNA damage (18), metabolic enzyme/protease (18), protein tyrosine kinase/RTK (17), JAK/STAT signaling (14) and membrane transporter (13), substantially inhibit *in vitro* growth of Mtb mc^2^ 6206 (Fig. 1a).

**Figure 1.**
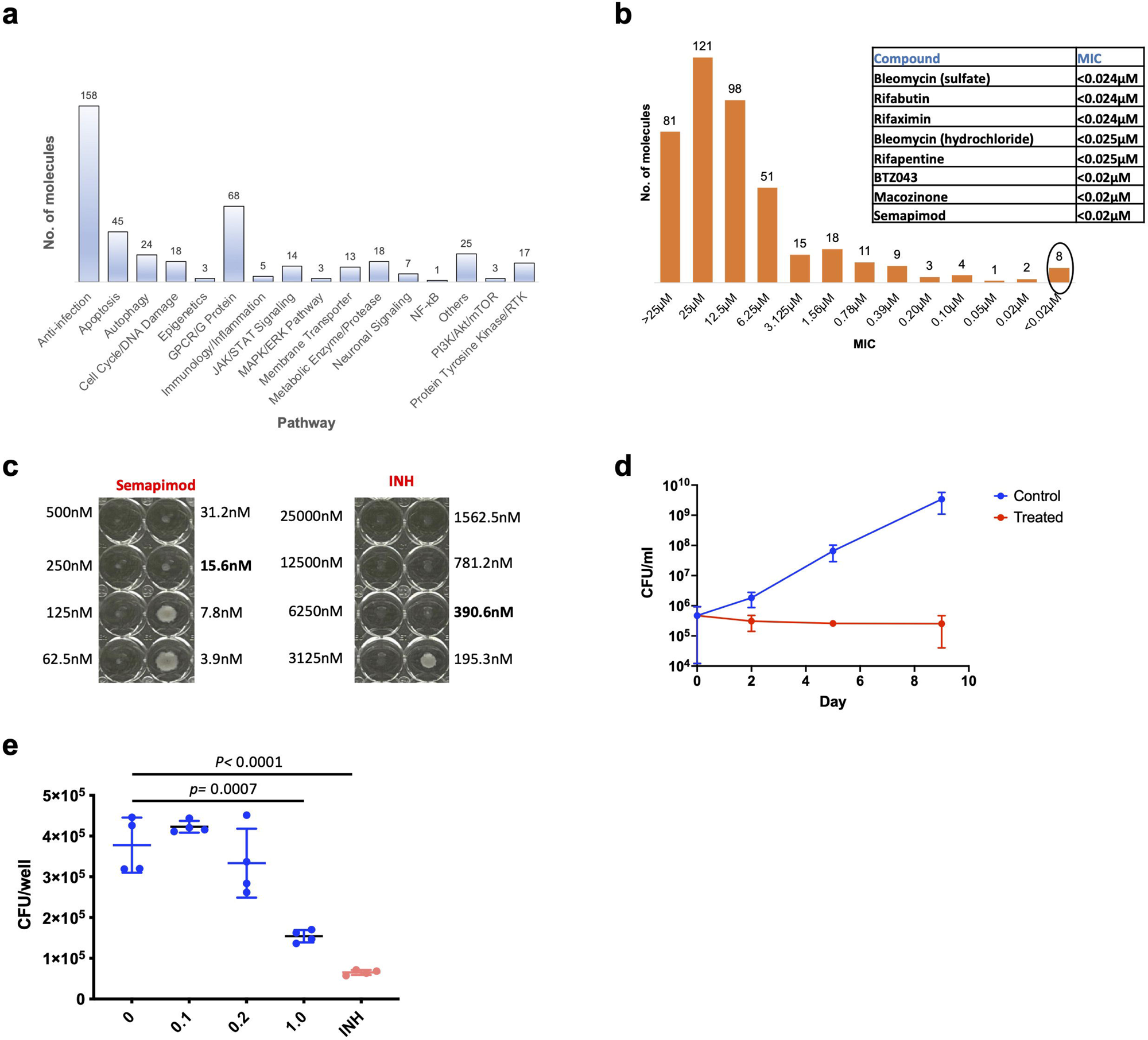
Screening of FDA-approved library of molecules against Mtb mc^2^ 6206 reveals growth-inhibitory effect of semapimod, an anti-inflammatory small molecule. **a**, Status of inhibitors showing activity against Mtb mc^2^ 6206. Bar-graph depicts Mtb mc^2^ 6206 inhibitors acting on different host metabolic pathways. **b**, MIC_90_ status of Mtb inhibitors. Bar-graph shows inhibitors with different MIC_90_ against Mtb mc^2^ 6206, *in vitro*. Molecules with <0.025µM MIC_90_ (encircled) are tabulated in the inset. **c**, Determination of minimum dose of semapimod for complete inhibition of Mtb mc^2^ 6206 growth by visual inspection. Isoniazid (INH) was used as a control in the 96-well plate-based assay. Drug concentration beyond which no visual growth is observed, is written in bold. **d**, Effect of semapimod on the *in vitro* growth of Mtb mc^2^ 6206. Shown is the CFU enumeration of drug-treated and untreated (control) bacteria at the indicated time points. **e**, Effect of semapimod on the intracellular proliferation of Mtb mc^2^ 6206. Intracellular growth of Mtb mc^2^ 6206 in the THP1-derived macrophages was examined after 6 days of infection in the absence (UT) or the presence of 0.10-1.0µM semapimod. Treatment with 2.5µM INH was used as control. A significant reduction in CFU counts of Mtb mc^2^ 6206 is observed in the presence of semapimod under *in vitro* culture conditions (**d**) as well as during intracellular (*p*< 0.005) growth (**e**). Data represent mean±s.d of at least n=2 replicates in d and n=4 replicates in **e**. Data in **c** are representative of n=2 independent experiments. *p* values in **e** were obtained after comparison of CFUs between the UT and semapimod-treated samples, as described in Materials and Methods.

Next, we determined minimum concentration of newly identified anti-mycobacterial compounds at which ∼90% growth is suppressed (MIC_90_), by using plate-based resazurin assay. Our results reveal 300 molecules showing an MIC_90_ of ≥12.5 µM, whereas remaining 122 molecules exhibit MIC_90_ values ranging from 6.25 to ≤0.025µM (Fig. 1b). Surprisingly, out of the eight molecules which inhibit growth at <0.025µM, we find an anti-inflammatory molecule namely semapimod which demonstrates anti-microbial activity against Mtb mc^2^ 6206. Notably, semapimod is an inhibitor of proinflammatory cytokine production, which suppresses the production of TNF-α, IL-1β, and IL-6^17, 18, 19, 20^, and its anti-mycobacterial activity has never been reported to the best of our knowledge.

### Semapimod shows bactericidal activity against Mtb mc^2^ 6206 under extra- and intracellular growth conditions

To corroborate the findings from initial screen, we tested the effect of the fresh batch of semapimod against Mtb mc^2^ 6206, which reveals complete inhibition of mycobacterial growth at ∼15nM (Fig. 1c). The colony forming unit (CFU) estimation further shows a bactericidal activity of this molecule which causes 88% reduction of bacterial viability on day 2 and >99% reduction after 5 days of incubation (Fig. 1d). In addition to its effect on *in vitro* growth, we also find a substantial reduction in viable bacterial counts during macrophage infection. Incubation of Mtb mc^2^ 6206-infected THP1 macrophages with different doses of semapimod reveals 60% reduction (*p* <0.005) in intracellular CFU counts after 6 days of incubation with 1µM inhibitor (Fig. 1e). Notably, no cytotoxic effect was observed at this concentration against THP1, thus ruling out the possibility of cell lysis by semapimod.

### Global transcriptional profile of Mtb in response to semapimod treatment

To gain mechanistic insights into the effect of semapimod on Mtb mc^2^ 6206, we analysed the global transcriptional profile of the bacterium in response to this drug. Bacterial cultures in the logarithmic growth phase (OD600 of ∼0.4) were treated with 50nM semapimod, equivalent to ∼3x MIC, and total RNAs were harvested from semapimod-treated and untreated bacteria after 24 h of incubation, as described in the Materials and Methods. RNA samples from three biological replicates were subsequently used in RNA sequencing (RNASeq), which reveals differential expression of 482 genes by ≥2.0-fold (false discovery rate (FDR)-adjusted *p* value of ≤ 0.01). While 251 genes are downregulated by semapimod treatment, expression of 231 genes is induced (Supplementary Dataset 1 and Fig. 2a). Remarkably, the effect of semapimod was consistent across three biological replicates (Fig. 2b). Genes, prominently affected by semapimod, primarily belong to intermediary metabolism and respiration (n=124), cell-wall and cell processes (n=98), lipid metabolism (n=50) and those encoding for regulatory proteins (n=31) (Fig. 2c). Expression of some of the representative genes was also validated by qRT-PCR using gene-specific primers, which shows a similar pattern of expression (Supplementary Fig. 2), thus confirming the specificity of the RNASeq data.

**Figure 2.**
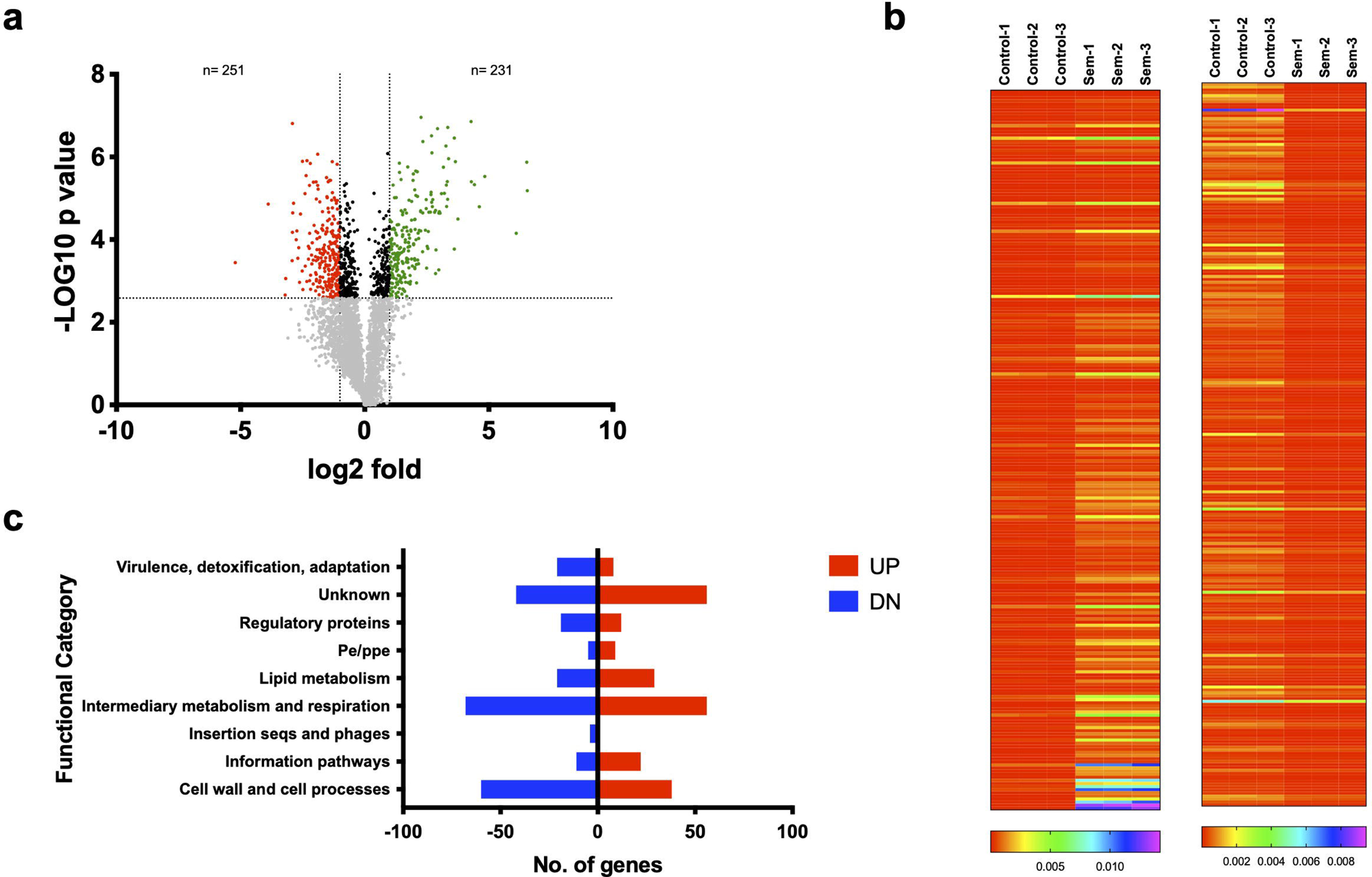
Effect of semapimod treatment on the expression profile of Mtb mc^2^ 6206 transcripts. **a**, Volcano plot of differentially expressed genes in semapimod-treated Mtb mc^2^ 6206. The plot shows distribution of genes that are differentially expressed *via* log2 fold-change and the –log *p* values. Broken vertical lines represent the cutoff of ≥1.0 log2 fold-change, and horizontal line represents the cutoff of >2.583 –log *p* values. Genes below the - log *p* cutoff are represented by grey dots. Downregulated genes are represented by red dots, and those showing upregulation in response to semapimod treatment are shown with green dots. **b**, Status of differentially accumulated transcripts. Heatmap representation of transcripts showing accumulation (left) or suppression (right) upon exposure to semapimod across three biological replicates. **c**, Functional categorization of differentially regulated genes. The butterfly chart shows distribution pattern of differentially regulated genes according to their function, as classified in the Mycobrowser database (https://mycobrowser.epfl.ch/genes/). Mean fold-change values from n=3 biological replicates are shown in **a**.

Our results show that genes involved in oxidative phosphorylation such as *atpF*, *atpH*, *atpG*, *atpD*, *cydB*, *cydA*, *Rv1812c* and *qcrC* as well as several *nu*o operon genes (*nuoB*, *nuoC*, *nuoD*, *nuoE*, *nuoF*, *nuoG*, *nuoI*, *nuoJ*, *nuoL*, *nuoM* and *nuoN*) are downregulated in the drug-treated bacteria. Not only expression level of these respiratory genes is perturbed, but also a significant reduction in the cellular ATP levels is observed upon semapimod treatment of Mtb mc^2^ 6206 (Supplementary Fig. 3). Furthermore, we find several genes that encode for subunits of RNA polymerase (*rpoA* and *rpoB*) and ribosome (*rplJ*, *rplL*, *rplC*, *rplD*, *rplW*, *rplB*, *rpmC*, *rpsN1*, *rpsH*, *rplF*, *rplR*, *rpmD*, *rplT* and *rpsD*) exhibiting distinct upregulation in response to the drug treatment, which indicate perturbation of transcription and translation machineries in the drug-treated bacteria. Notably, semapimod treatment also interferes with expression of a number of genes associated with metabolism of amino acids such as branched-chain amino acids (*ilvC*, *ilvB1*, *ilvN*, *leuA*, *bkdB*, *bkdA*, *Rv2499c*, *accA1* and *accD1*), cysteine (*csd*, *cysM*, *cysO*, *cysE*, *cysK1* and *sahH*), lysine (*pcd*, *lat*, *lysA*), methionine (*metZ*, *metK* and *metH*), threonine (*thrB*, *thrC* and *thrA*), glutamate (*gdh*), glutamine (*glnA1*) and tryptophan (*trpA*).

Among the cell-wall and cell processes functional category, we find a substantial reduction in the expression of genes involved in peptidoglycan biosynthesis (*pbpA*, *ponA1*, *murI*, *dacB1*), cell-wall arabinogalactan linker formation (*wbbL1*), cell-wall arabinan biosynthesis (*embB* and *aftB*), cell-wall LPS biosynthesis (*Rv0225*, *gca*, *gmhA*, *gmhB* and *hddA*), cell division (*Rv1708*, *scpA*, *ftsZ* and *gid*) and transport across the bacterial cell membrane (*ctpB*, *glnH*, *ctpE*, *pstA1*, *pstA2*, *narK2*, *Rv1747*, *ansP1*, *fecB* and *Rv3200c*).

Another category of genes exhibiting notable change in expression are those involved in the biosynthesis of lipids such as cell-wall mycolic acid (*accD6*, *fabD*, *kasA*, *kasB*, *fabG4*, *accD4*, *umaA*, and *cmaA2*), methyl-branched lipids (*ppsB*, *ppsC*, *ppsD*, *mas*, *fadD26*, *fadD29*, *tesB1*, *Rv2953*, *papA1*, *pks2*, *mmpL8*, *fadD23*, *pks3*, *pks4*, *papA3* and *fadD21*), phospholipids (*cdh*) and triglycerides (*desA3*). In addition, several genes controlling lipid degradation (*fadE5*, *fadE35*, *fadD12*, *fadE19*, *tesB2*, *fadE21*, *fadE24*, *fadB2*, *fadD36*, *fadE15* and *fadD9*) and transport (*lprK*, *mce1F*, *mce4F*, *mce4D*, *lprN*, and *lucA*) are also modulated in the drug-treated bacteria. Noteworthy to mention, a marked reduction in expression is observed for genes involved in synthesis of succinyl-CoA *via* vitamin B12-dependent methylmalonyl pathway (MMP; *accA3*, *mutA*, *mutB*, *cobB* and *cobO*), whereas those associated with methylcitrate cycle (MCC; *prpD*, *prpC*, *icl1* and *pks11*) exhibit significant upregulation, together indicating perturbation of acetyl CoA levels in semapimod-treated Mtb mc^2^ 6206.

Furthermore, we find that semapimod treatment modulates expression of genes encoding a variety of regulatory proteins such as kinases (*pknA*, *pknG* and *pknD*), WhiB-family transcription regulators (*whiB2*, *whiB3* and *whiB7*), hypoxia regulators (*Rv0081*, *dosS* and *dosR*), regulators of lipid transport (*Rv0302* and *mce1R*), a regulator of MCC genes (*regX3*) and a leucine-responsive transcription regulator (LrpA).

### Semapimod treatment interferes with L-leucine uptake in Mtb

To validate the effect of semapimod on the virulent Mtb, we sought to determine its MIC against Mtb H_37_Rv. Surprisingly, the drug was found ineffective against virulent Mtb strain. Similar to virulent Mtb strain, no effect was observed against other mycobacterial species such as *M. bovis* BCG, *M. abscessus* and *M. smegmatis* (Fig. 3a). These results pointed out the effect of semapimod on uptake of leucine and/or pantothenate that are indispensable for growth of the auxotrophic strain. Altered expression of a set of LrpA-regulated genes^21, 22, 23^ by semapimod treatment indicates perturbation of leucine uptake. To test this hypothesis, we cultured Mtb mc^2^ 6206 in the absence of pantothenate and L-leucine (PL-), pantothenate (P-) or L-leucine (L-) for 24 h followed by analysis of the expression of some of the leucine-responsive genes such as *lat*, *leuA*, *lrpA* and *icl1* as surrogate markers. Interestingly, all of these genes exhibit marked increase in expression under PL- and L-, but not under P-conditions (Supplementary Fig. 4). A similar expression profile of these genes is observed in semapimod-treated bacteria by RNASeq, thus suggesting the interference of L-leucine uptake. To corroborate these findings, the intracellular leucine concentrations were estimated in the untreated and semapimod-treated Mtb mc^2^ 6206 by mass-spectrometry. Our results show ∼45% reduction (p <0.05) in the leucine levels in bacteria exposed to semapimod (Fig. 3b). Noteworthy to mention, levels of another BCAA, valine or an unrelated amino acid, L-proline remain unchanged in Mtb mc^2^ 6206 upon semapimod treatment, which corroborates effect of the drug specifically on L-leucine uptake (Supplementary Fig. 5). Next, we examined the effect of semapimod against Mtb mc^2^ 6206, expressing *leuC-leuD* and *panC-panD*, respectively, under a constitutive promoter. Remarkably, expression of *leuC-leuD*, and not of *panC-panD* genes, alleviates the growth-inhibitory effect of semapimod (Fig. 3c-e). Mtb mc^2^ 6206::*leuCD* exhibits comparable growth in the presence and the absence of semapimod (Fig. 3c). In addition, Mtb mc^2^ 6206::*leuCD* exhibits resistance to killing by as high as 3.37 µM (>200-fold excess of MIC) semapimod (Fig. 3d-e). In contrast, Mtb mc^2^ 6206::*panCD* exhibits a similar level of inhibition by semapimod, as observed with the Mtb mc^2^ 6206. Overall, these results substantiate that semapimod inhibits L-leucine uptake in Mtb.

**Figure 3.**
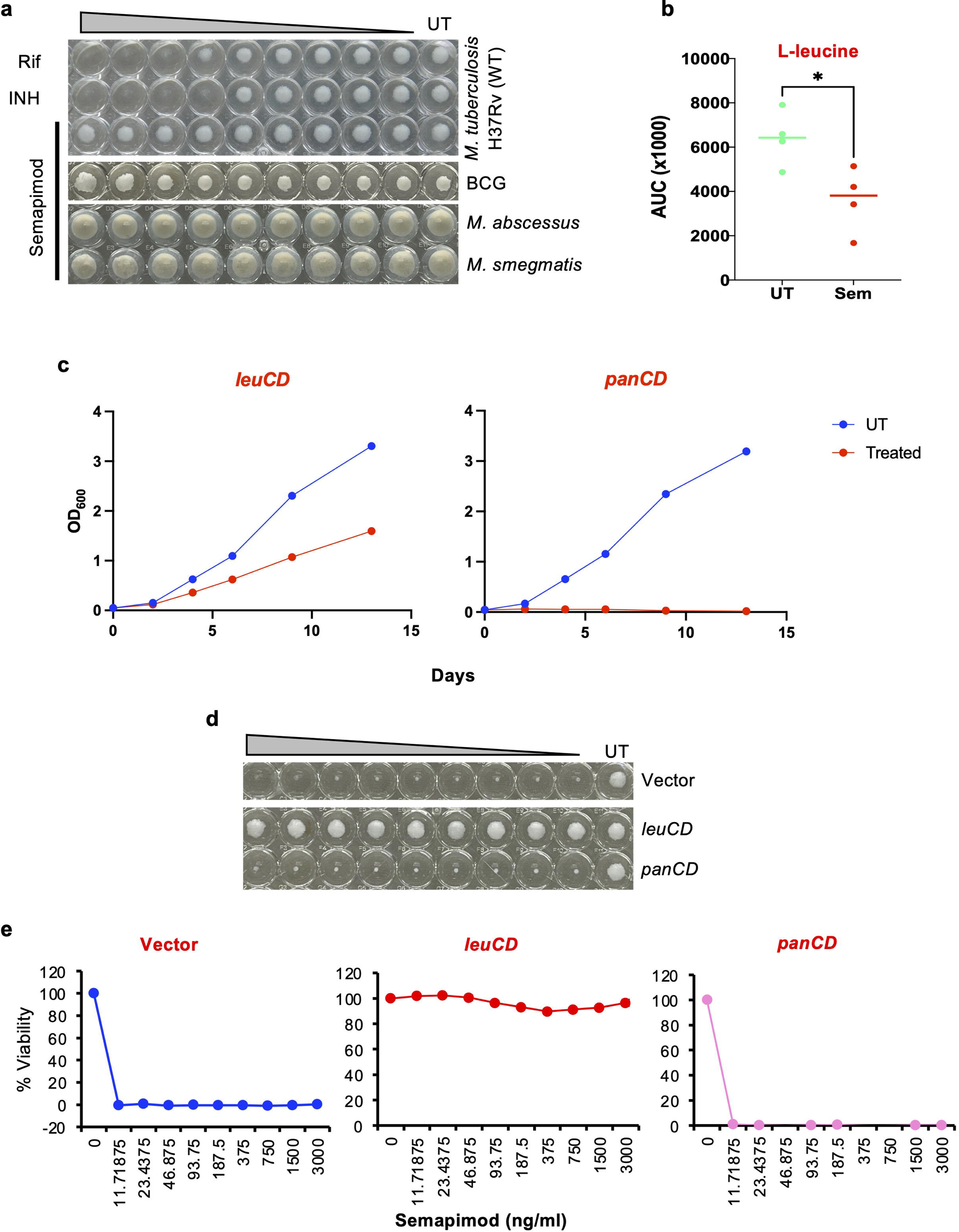
Semapimod treatment perturbs L-leucine uptake in Mtb mc2 6206. **a**, Effect of semapimod on the *in vitro* growth of different mycobacterial species. Growth inhibitory effect of semapimod was assessed against slow-growing Mtb H_37_Rv and *M. bovis* BCG, and fast-growing *M. abscessus* and *M. smegmatis*, respectively, by visual inspection using 96-well plate-based assay. Rifampicin (Rif) and INH were used as control drugs against Mtb H_37_Rv. **b,** Estimation of intracellular leucine in Mtb mc^2^ 6206. Intracellular leucine was estimated in untreated (UT) and semapimod-treated (Sem) bacteria, after 24 hours of drug treatment, by mass spectrometry as described in Materials and Methods. **c**–**e**, Effect of *leuC-leuD* and *panC-panD* expression in Mtb mc^2^ 6206 on bacterial killing by semapimod. Time- (**c**) and dose- (**d**–**e**) dependent growth kinetics reveals loss of bactericidal effect of semapimod against Mtb mc^2^ 6206 by *leuC-leuD*, and not by *panC-panD* expression. Percent viability in **e** was calculated with respect to untreated (UT) cultures after 2 weeks of exposure to different drug concentrations, using 96-well plate-based method as described in Materials and Methods. Data in **a** and **c**–**d** are representative of n=2 independent experiments. Mean±s.d values from n=4 replicates are shown in **b.**

### Generation and characterization of semapimod resistant strain of Mtb mc^2^ 6206

Semapimod-mediated inhibition of L-leucine uptake suggests the presence of a committed L-leucine transport machinery in mycobacteria. While a number of putative L-leucine transporter genes are found in *M. smegmatis* (Supplementary Fig. 6), none of their orthologues are present in Mtb.

To gain insights into the potential mechanism of L-leucine transport in Mtb, we generated the semapimod resistant (Sem^R^) strain of Mtb mc^2^ 6206 by repeated passaging in the presence of drug (Fig. 4a). *In vitro* growth analysis reveals a significantly increased growth of the two independent Sem^R^ strains (Fig. 4b). While, the wild-type bacteria do not grow beyond OD_600_ of 2.0, both the Sem^R^ strains achieve an OD_600_ of ∼4.0 after 10 days of culturing in 7H9-OADS medium supplemented with L-leucine and pantothenate (Fig. 4b). Remarkably, intracellular L-leucine, but not the other BCAA such as valine, is significantly higher in the Sem^R^ strain when compared with the WT bacteria (Supplementary Fig. 7). The semapimod resistance does not lead to any change in susceptibility to standard TB drugs such as rifampicin or isoniazid (Supplementary Fig. 8), however, a marked increase in susceptibility to vancomycin is observed in the Sem^R^ bacteria. While, vancomycin inhibits Sem^R^ with the IC_50_ of ∼7µg/ml, no effect is observed for this antibiotic against the wild-type Mtb mc^2^ 6206 (Fig. 4c). It has been reported earlier that bacteria lacking PDIM grow faster and exhibit more susceptibility to vancomycin than the wild-type Mtb with intact PDIM which provides barrier to nutrients as well as large molecules such as vancomycin^24^. Interestingly, whole genome sequence analysis of the Sem^R^ strain reveals Ile896Asn mutation in PpsB, involved in PDIM synthesis (Fig. 4d). In addition, three other genes show altered sequence in Sem^R^ namely, *mce1D*, *Rv3837c* and *ppe60*. While *mce1D* accumulates a nonsense mutation yielding a stop codon at 225^th^ position, sequence alteration in *Rv3837c* is at the extreme 3’ end causing Thr ➔ Ile substitution at the 228^th^ position of the 332 amino acid long polypeptide. Furthermore, multiple mutations were observed exclusively in *ppe60* (Supplementary Fig. 9), despite the presence of two other paralogues of this gene (*ppe18* and *ppe19*), suggesting random accumulation of mutations in *ppe60* during passaging. These observations led us to investigate the association of *ppsB, mce1D* and *ppe60* in semapimod resistance, whereas, the possibility of involvement of *Rv3837c* is unlikely.

**Figure 4.**
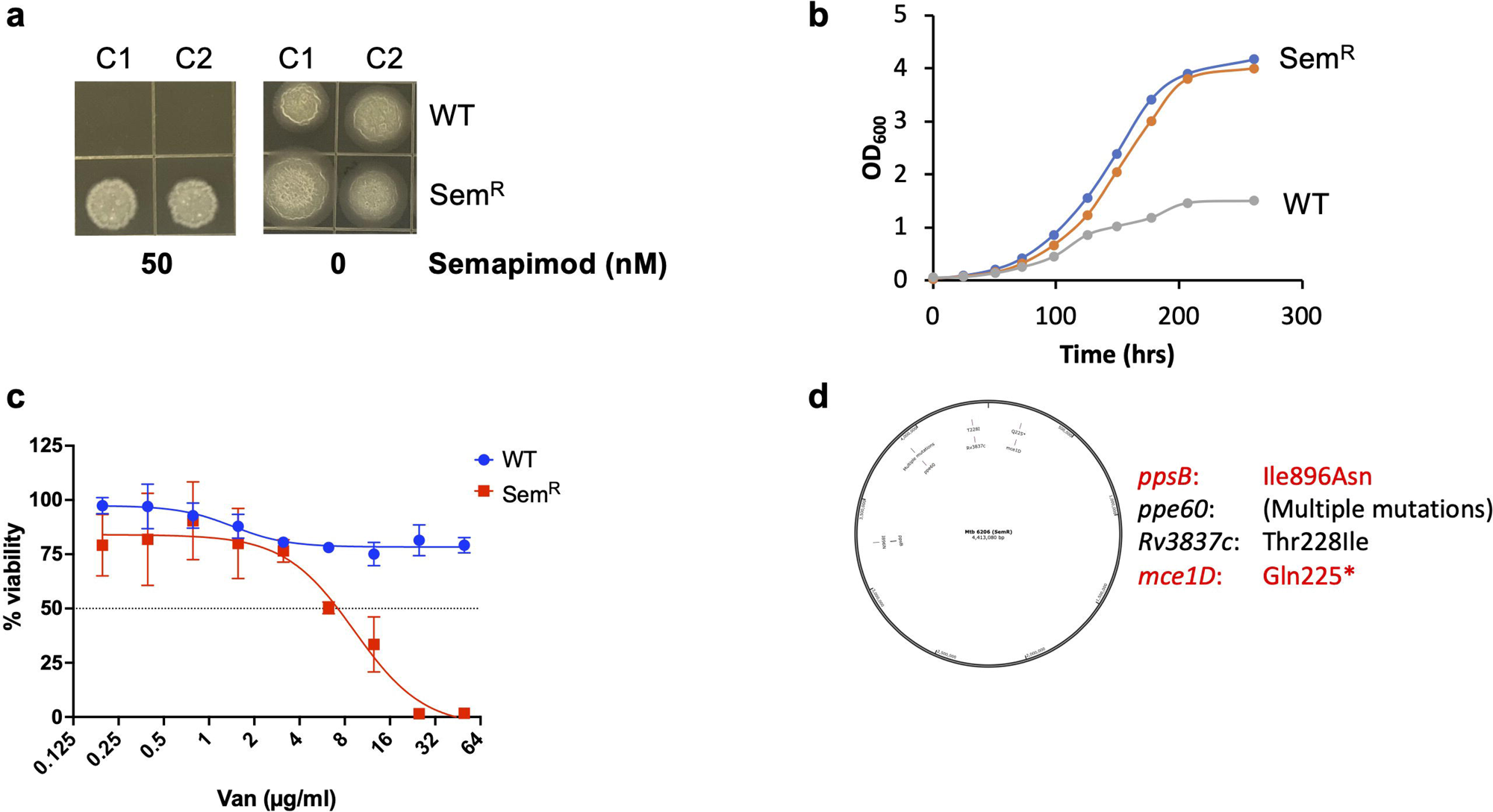
Generation and characterization of semapimod-resistant strain of Mtb mc^2^ 6206. **a**, Propagation of putative semapimod-resistant (Sem^R^) strain, but not the wild-type (WT) Mtb mc^2^ 6206 in the presence of 50nM semapimod confirms drug-resistance in Sem^R^. In contrast, both the strains exhibit substantial growth on 7H11-PLO agar plate in the absence of drug. Shown is the growth pattern of two different colonies– C1 and C2 of the respective strains. **b**, *In vitro* growth analysis of the WT and the Sem^R^ strains of Mtb mc^2^ 6206. Comparative analysis of OD_600_ at different time points reveals substantially increased growth of Sem^R^ strain compared to WT under *in vitro* culture conditions. **c**, *In vitro* susceptibility of the WT and the Sem^R^ strains of Mtb mc^2^ 6206 to vancomycin. Viability of WT and Sem^R^ Mtb strains was determined in the presence of vancomycin by 96-well plate-based assay, as described in Materials and Methods. Percent viability was calculated with respect to the untreated (UT) cultures after 2 weeks of exposure to different concentrations of antibiotic. Results show a substantial increase in susceptibility of Sem^R^ to vancomycin compared to WT. **d**, Whole genome map analysis of Sem^R^. Shown is the genome map of Sem^R^ highlighting positions of genes undergoing substitutions, when compared with WT Mtb mc^2^ 6206. Mutations leading to corresponding changes at the amino acid level in the respective protein are indicated alongside. Suspect genes presumably involved in providing semapimod resistance, are marked in red fonts. Data are representative of at least n=2 independent experiments in **a**–**b**. Mean±s.d values from n=3 biological replicates are shown in **c.**

### Overexpression of *ppsB* alters susceptibility of semapimod^R^ strain to drugs

To further identify genes that might contribute to semapimod resistance, we overexpressed *ppsB, mce1D* and *ppe60* genes in the wild-type and Sem^R^ strains of Mtb mc^2^ 6206 under the constitutive promote. Notably, susceptibility to vancomycin and semapimod was completely reversed in Sem^R^ strain upon overexpression of *ppsB*, whereas no effect of *ppsB* overexpression is observed in the wild-type bacteria (Fig. 5). Sem^R^ strain, which is susceptible to vancomycin, becomes resistant akin to the wild-type strain by overexpression of *ppsB* (Fig. 5a). Similarly, susceptibility to semapimod is restored in Sem^R^ by overexpressing *ppsB* (Fig. 5b). In contrast, no effect of *mce1D* or *ppe60* overexpression is observed on bacterial sensitivity to semapimod (Supplementary Figs. 10 and 11).

**Figure 5.**
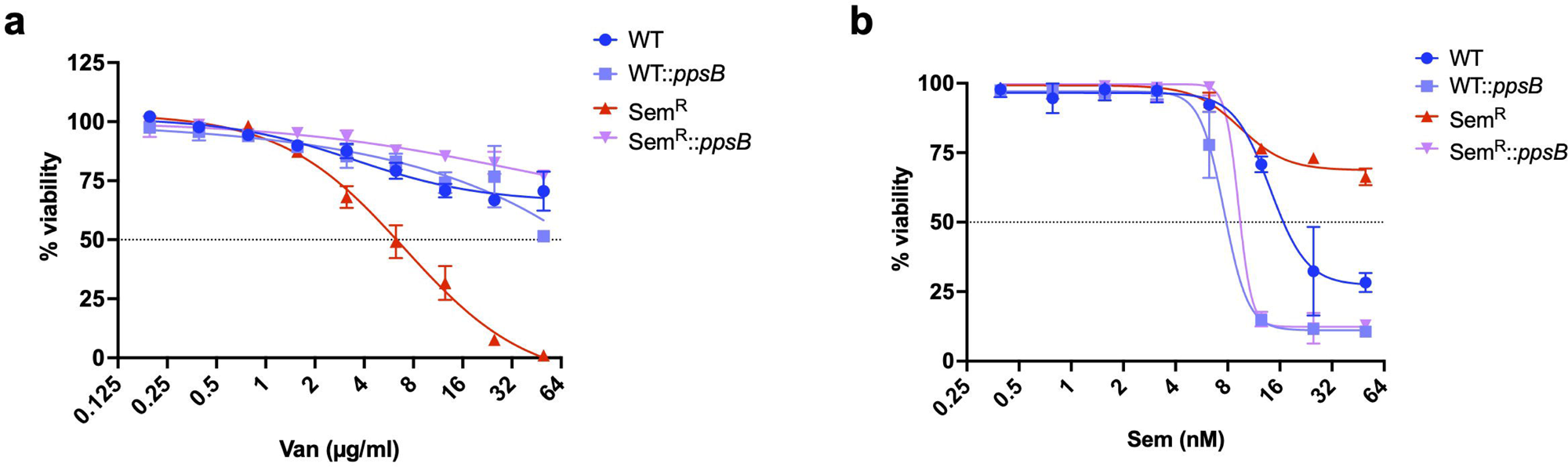
Overexpression of *ppsB* alters susceptibility of Sem^R^ strain of Mtb mc^2^ 6206 to semapimod and vancomycin. **a**–**b**, *In vitro* susceptibility of different strains of Mtb mc^2^ 6206 to vancomycin (**a**) and semapimod (**b**). Viability of WT and Sem^R^ Mtb strains in the presence of drugs was compared with those constitutively expressing *ppsB* by 96-well plate-based assay, as described in Materials and Methods. Percent viability was calculated with respect to UT cultures after 2 weeks of exposure to different concentrations of inhibitors. Remarkably, response of Sem^R^ to both semapimod and vancomycin is reversed upon overexpression of *ppsB*. Contrarily, *ppsB* expression in WT does not affect bacterial sensitivity to either of these drugs, thus indicating a specific effect of the PDIM biosynthesis gene in Sem^R^. Mean values from n=3 biological replicates are shown in **a**–**b**.

### Semapimod targets PpsB of *M. tuberculosis*

The above results strongly suggest that the cell-wall PDIM biosynthesis protein PpsB is involved in semapimod-mediated interference of L-leucine uptake in Mtb. To corroborate our findings, we sought to examine if PpsB of Mtb is directly targeted by semapimod. The 4617bp *ppsB* open reading frame of Mtb was cloned in an *E. coli* expression plasmid, pET28, for overexpression and purification of 6x His-tagged PpsB of Mtb. Analysis of purified protein by SDS-PAGE reveals near-homogeneous preparation of the protein migrating at its predicted molecular mass of ∼170kDa (Supplementary Fig. 12). The purified 6x His-PpsB was then subjected to interaction with different concentrations of semapimod using Biolayer Interferometry-Octet system (Sartorius), as mentioned in the Materials and Methods. The results, presented in Fig. 6a, clearly demonstrate a dose-dependent interaction of semapimod with PpsB, immobilized on AR2G sensor. Further analysis of binding kinetics reveals that semapimod strongly binds to PpsB with a dissociation constant (*K_d_*) of 110nM. As a control, interaction of semapimod was also analysed with the purified Ppe60, which fails to exhibit any binding.

**Figure 6.**
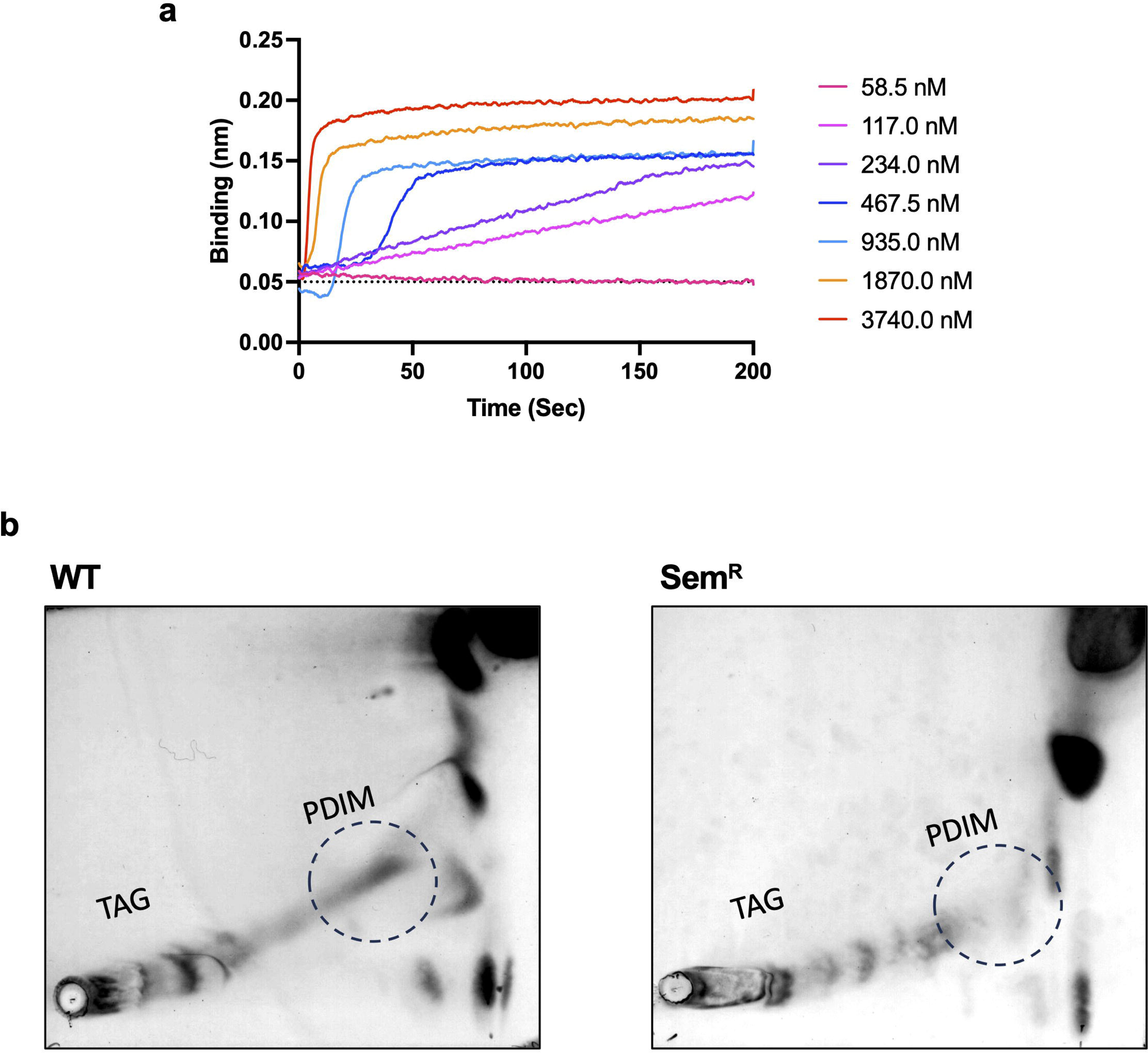
Semapimod targets PDIM biosynthesis protein, PpsB. a, Analysis of semapimod-PpsB interaction by BLI-Octet. Interaction of semapimod with Mtb PpsB was examined by optical interference–based biolayer interferometry from the Octet system (ForteBIO). Briefly, dialyzed 6xHis-PpsB was immobilized onto AR2G sensor up to a level of 2.1 nm. Binding was observed at the indicated concentrations of ligand (semapimod) to acquire differential graded response. Binding constant was calculated as per the standard steps, described in Materials and Methods. **b**, Analysis of cell-wall apolar lipids in the WT and Sem^R^ strains of Mtb mc^2^ 6206. Cell-wall apolar lipids were extracted and analyzed by two-dimensional TLC as described in Materials and Methods, which reveals significant reduction in PDIMs of Sem^R^ when compared with the WT strain. Contrary to PDIMs, no change is observed in the triacyl glycerol (TAG) levels between the two strains. Position of PDIMs is marked by broken circle in both the images for clarity.

### PDIM is compromised in Sem^R^ strain

As hitherto mentioned, PDIM has been found to act as a barrier for uptake of nutrients and large-size inhibitors. Moreover, our results clearly establish that semapimod targets PDIM biosynthesis protein PpsB by direct binding. Based on our findings, we speculate that semapimod resistance is acquired in Mtb owing to disruption in the cell-wall PDIM, which alleviates the regulation of L-leucine uptake. To test our hypothesis, we analysed the cell-wall lipids of the wild-type and Sem^R^ strains of Mtb mc^2^ 6206. Both mycolic acids and PDIM were extracted from these strains and analysed by thin layer chromatography, as described in the Materials and Methods. While no difference in the profiles of mycolic acid methyl esters or the tri-acyl glycerol (TAG) was noticed between the two strains, the Sem^R^ exhibits a marked reduction in PDIM (Supplementary Fig. 13 and Fig. 6b).

### Intracellular survival of Mtb H_37_Rv pathogen in the organelles of infected BalB/C mice is decreased upon semapimod treatment

Although, endogenous leucine biosynthesis is extremely essential for bacterial survival and virulence, there is no study describing underlying mechanism of L-leucine uptake from extracellular environment and its importance in mycobacterial virulence during animal infections. To examine if L-leucine uptake is critical for intracellular survival of the pathogenic Mtb H_37_Rv strain with intact endogenous leucine biosynthesis, we tested the effect of semapimod treatment on intracellular burden of Mtb H_37_Rv during mouse infection. Infection was performed via aerosol route as described in the Materials and Methods. After 21 days of infection, a group of 3 mice were treated with 5mg/kg dose of semapimod, whereas another group of 3 mice were treated with vehicle (1x phosphate buffered saline) by daily oral gavaging for 4 weeks, typically as shown in the schematic. Bacterial load was examined in lungs and spleen of animals from both the groups by colony forming unit (CFU) estimation (Fig. 7a). Examination of the gross lung pathology reveals a large number of granulomatous nodules in the untreated infected group, whereas semapimod treatment led to a significant reduction in the disease pathology (Fig. 7b). Remarkably, semapimod treatment caused ∼82% and ∼85% reduction in CFU counts in lungs and spleen, respectively, after 4 weeks of semapimod treatment (Fig. 7c and d). Noteworthy to mention, *in vitro* susceptibility to vancomycin remained unaffected in bacteria retrieved from lungs of the untreated and semapimod-treated mice (Supplementary Fig. 14), thus indicating comparable levels of cell wall PDIMs, one of the virulence factors, in both untreated and semapimod-exposed Mtb H_37_Rv.

**Figure 7.**
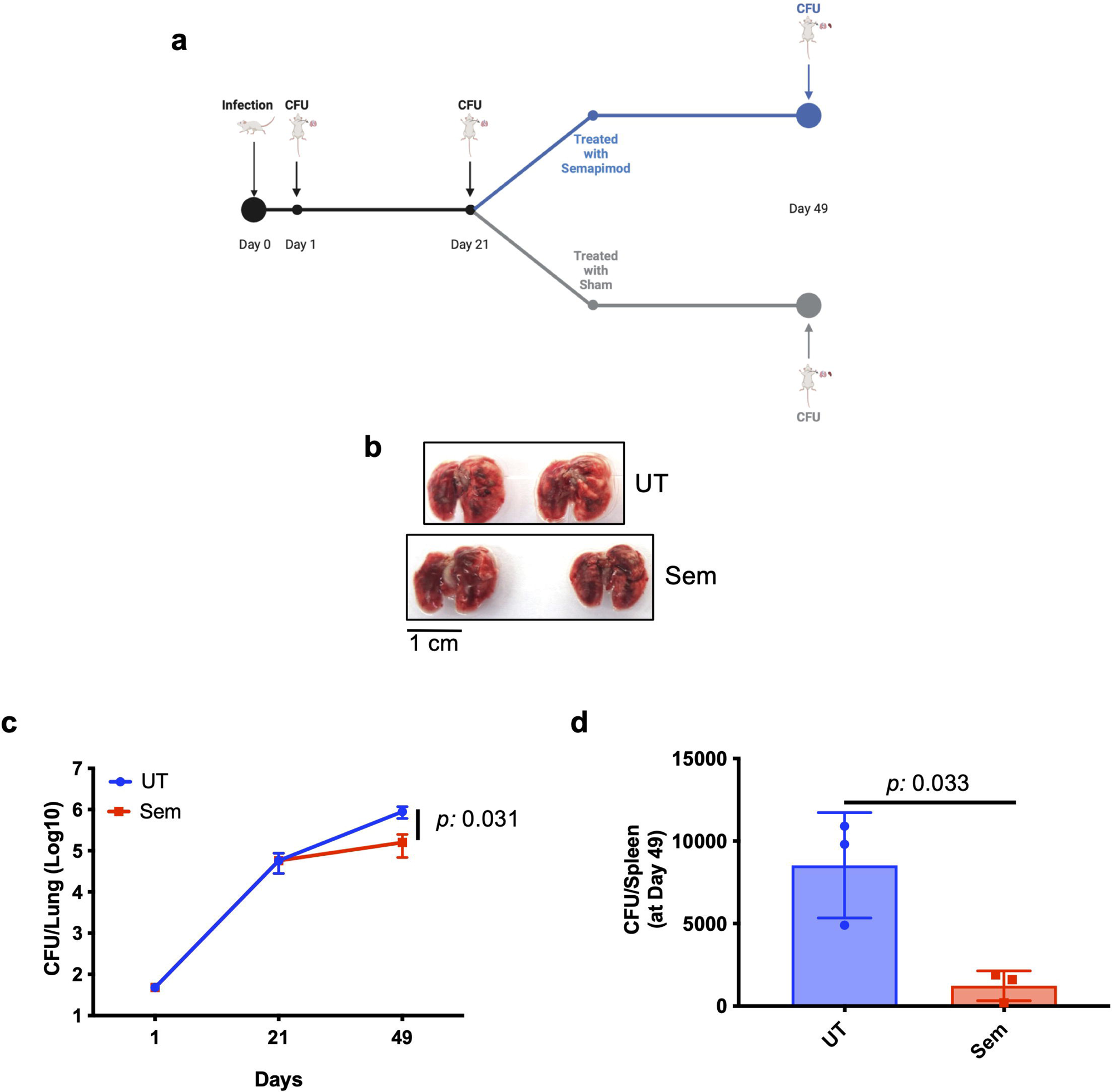
Effect of semapimod treatment on survival of Mtb H37Rv in the organelles of infected BALB/c mice. **a**, Schematic of mouse infection. Infection was performed by aerosol route with the virulent Mtb H_37_Rv strain. After 21 days of infection, mice were divided into two groups: one receiving only 5% sucrose (sham) and others receiving 5mg/kg semapimod prepared in 5% sucrose. Intracellular bacterial load was determined by CFU plating of lung homogenates at days 1, 21 and 49, and in spleen homogenates prepared on day 49. **b**, Gross pathology of lungs. Images of lungs obtained from both the sham- and semapimod-treated groups of mice after 28 days of treatment (i.e., day 49 post-infection) are presented. Scale bar is shown for size reference. **c**–**d**, Effect of semapimod treatment on intracellular survival of Mtb H_37_Rv. Intracellular survival was determined by estimating the bacterial burden in lungs at the respective time points (**c**), and in spleen at day 49 post-infection (**d**), by CFU enumeration. Data represent mean ± s.d. values from n=3 animals in **c**–**d.** *p* values in **c** and **d** were obtained after comparison CFUs between the two groups, as described in Materials and Methods.

Overall, these results provide first evidence of the effect of a repurposed compound obstructing the *in vitro* L-leucine uptake on Mtb virulence and TB pathogenesis during infection.

## Discussion

Conventional approaches of drug discovery involve screening of NCEs against either the validated ‘drug target’ (target-based screen) or the pathogen as a whole (phenotypic screen)^25^. The aim is to identify inhibitor(s) showing a novel MoA. However, any promising NCE faces a multitude of barriers to reach to market launch from initial development^26^. Repurposing offers the possibility of identifying new MoA of the existing drugs that are already approved for human use^27, 28^. Therefore, it bypasses the traditional drug discovery and development pipeline involving *de novo* synthesis of NCEs. Several non-antibiotic drugs that are currently approved or under development may inhibit microbial pathogens, and therefore can be explored as novel antimicrobials, either alone or in combination with the existing antibiotics^29, 30^.

The present study was designed with the aim to identify novel TB inhibitors using the FDA-approved repurposed library comprising of over 3600 molecules. Remarkably, in addition to the known anti-TB drugs, we found several novel hit molecules that were developed against non-communicable diseases such as cancer, cardiovascular, metabolic and neurological diseases (Fig. 1a-b) that can be explored as novel anti-TB drugs in future. Notably, one of the top hits, namely semapimod, which inhibited Mtb mc^2^ 6206 growth at less than 20nM concentration, has never been reported for antibiotic activity. We show that this drug, not only kills Mtb mc^2^ 6206 *in vitro*, but also significantly reduces bacterial growth during macrophage infection (Fig. 1c-e). Surprisingly, semapimod did not exhibit growth inhibition of other mycobacteria including the pathogenic Mtb H37Rv (Fig. 3), which was used to develop the auxotroph by deletion of *leuC-leuD* and *panC-panD* genes. Remarkably, we demonstrate that growth inhibition by semapimod is due to blockage of L-leucine availability in Mtb mc^2^ 6206 (Fig. 3). We rule out the possibility of L-leucine quenching or modification by semapimod, as both the *leuC-leuD*-expressing and the Sem^R^ strains of Mtb mc^2^ 6206 continue to metabolize exogenous L-leucine in the presence of drug. Instead, our results suggest that semapimod restricts L-leucine uptake in Mtb. As a consequence, this drug is not effective against other mycobacterial species with the functional endogenous leucine biosynthesis pathway.

Cell membrane acts as a permeability barrier to foreign substances and selectively filters molecules that are essential for the cellular physiology. Transport of solutes involves three different classes of transport mechanisms, viz., passive diffusion, facilitated diffusion and active transport. Of these three different classes, amino acid transport fulfil the criteria of facilitated diffusion and active transport^31^. The two best characterized amino acid transport systems are the high-affinity periplasmic binding protein-(BP) dependent transport system for leucine, isoleucine, and valine (LIV-I and LIV-II) in *E. coli* and the high-affinity histidine transport system in *S. typhimurium* ^32, 33, 34, 35, 36^. Although, Mtb is able to synthesize all 20 amino acids^37, 38, 39^, however, to ensure constant supply of amino acids under a wide variety of environments, the TB pathogen must be equipped for their transport. Genome analysis of the fast-growing soil organism, *M. smegmatis* reveals 15 genes spread across three different loci that are putatively involved in BCAA transport (Supplementary Fig. 6). However, homologues of none of these are found in Mtb. Although, intracellular Mtb exploits multiple host-derived amino acids as nitrogen sources including BCAAs during growth in macrophages^40, 41^, thus far, it remains unclear how amino acids are transported across the cell envelope of Mtb. To the best of our knowledge, this is the first study providing evidence for the presence of a functional transport system for a BCAA in Mtb. We show that semapimod, which is an anti-inflammatory drug, targets a polyketide synthase involved in the cell-wall PDIM biosynthesis, and selectively kills *in vitro* the leucine auxotroph by blocking the L-leucine uptake, possibly by modulating the PDIM architecture. The cell-wall PDIM regulates uptake of nutrients and provides barrier to large molecule inhibitors. PDIM tends to shed during culturing of Mtb in the presence of ionic detergents, thus providing better access to nutrients and large molecules such as vancomycin. As a consequence, Mtb strains lacking PDIM not only grow faster, but also become more susceptible to vancomycin^24^. Remarkably, the Sem^R^ strain which accumulates mutation in the PpsB near acetyltransferase domain, exhibits both the phenotypes, i.e., a dramatic increase in its susceptibility to vancomycin and faster growth rate compared to the wild-type Mtb, despite culturing in the presence of the non-ionic detergent. Loss of PDIM in the drug-resistant strain is further confirmed by TLC analysis of the PDIM, whereas other lipids such as triacyl glycerol (TAG) or mycolic acids remain unaltered (Fig. 6b and Supplementary Fig. 13). Most importantly, susceptibility to semapimod is restored in the Sem^R^ Mtb mc^2^ 6206 by overexpression of *ppsB*. Taken together, these results clearly demonstrate for the first time the involvement of the cell-wall PDIM in L-leucine uptake in Mtb (Fig. 8). At present, it is not understood how PDIM selectively facilitates movement of L-leucine, and whether other apolar hydrophobic molecules follow the same route for their transport across the mycobacterial cell membrane. Since, PDIM is an important virulence factor^42^, understanding its role in the transport of other metabolites including amino acids will shed important light on virulence strategies employed by Mtb.

**Fig. 8.**
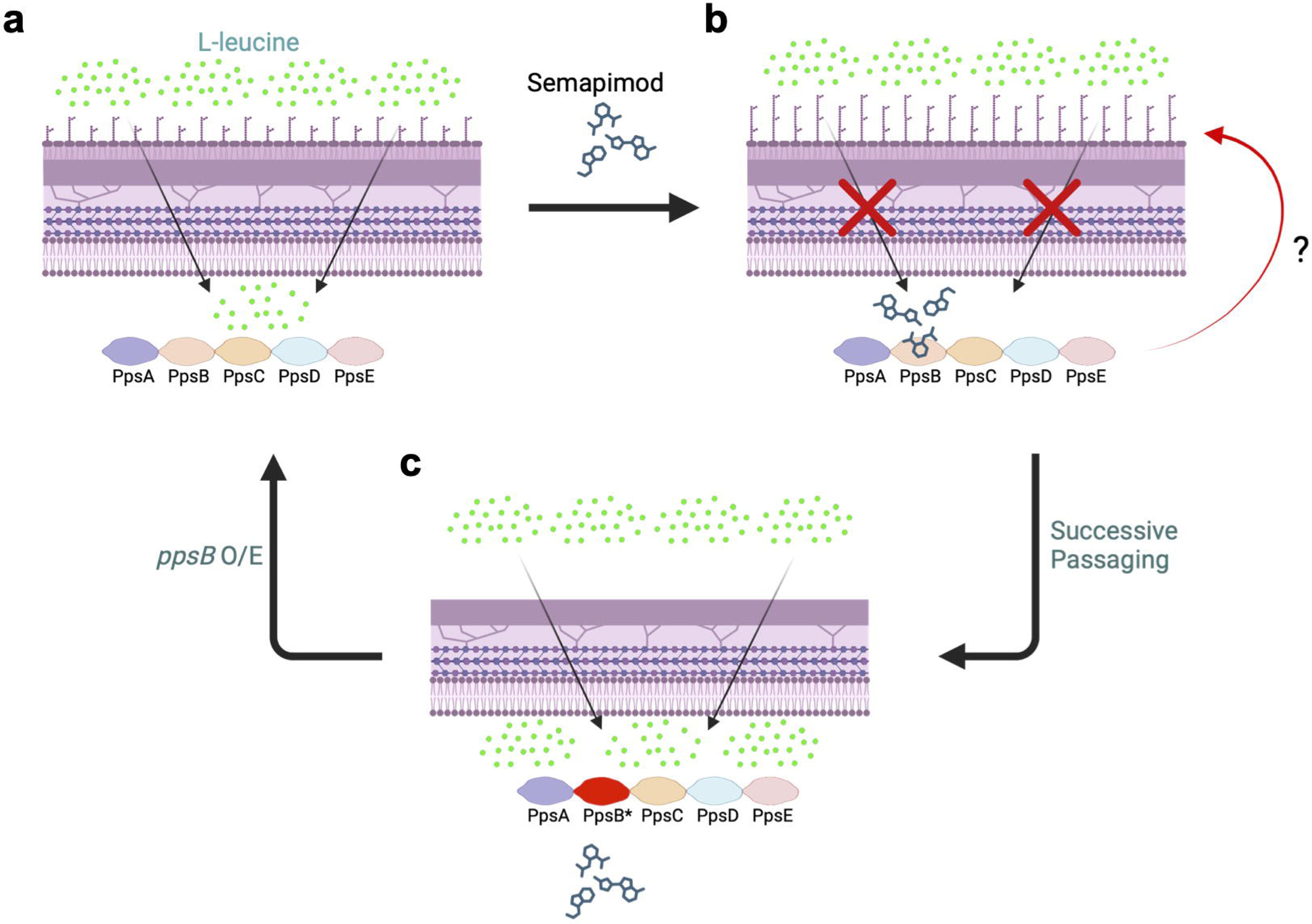
Proposed model describing the effect of semapimod on L-leucine uptake in Mtb mc^2^ 6206. **a**, Uptake of L-leucine is controlled by cell-wall PDIM. **b**, Semapimod targets PDIM biosynthesis protein (PpsB) causing altered PDIM profile which restricts L-leucine uptake in the auxotroph by an unknown mechanism (denoted by ‘?’), leading to death. **c**, L-leucine is freely accessible to Sem^R^ strain, owing to perturbation in the cell-wall PDIM. As a consequence, semapimod is ineffective against this strain. However, overexpression of wild-type *ppsB* in Sem^R^ reverts the bacterial susceptibility to semapimod. The illustration was created in BioRender. (Agarwal, N. (2025) https://BioRender.com/h04h035).

Semapimod treatment interferes with the expression of more than 10% of protein-coding genes of Mtb involved in a variety of functions. A careful analysis of the transcriptional profile further provides evidence corroborating the deviation of leucine levels in the cell by semapimod treatment. For instance, *lrpA*, which encodes for a transcriptional regulator of the leucine-responsive regulatory protein (Lrp) family^43^, is upregulated by 9.25-fold. It has been shown that binding of Mtb LrpA with the target promoters is influenced by amino acids and vitamins^43^. Not only LrpA, but also some of its regulons such as *lat* (94.01-fold), *whiB2* (0.33-fold), *pks2* (12.58-fold), and *gid* (0.37-fold) also show significant modulation.

Among other differentially expressed gene sets, the most notable ones are those involved in energy and lipid metabolism. Since, leucine is catalyzed into acetyl-CoA^44^, we hypothesize that in the presence of semapimod, level of acetyl-CoA is perturbed in the cell. Acetyl-CoA, being the precursor of energy metabolism, subsequently impacts the overall energy state, as evidenced by >80% reduction in ATP following drug treatment. In other bacteria such as Salmonella, L-leucine supplementation stimulates the TCA cycle, causing increased intracellular NADH and ATP concentration^45^, which further substantiates the contribution of intracellular leucine in energy production in mycobacteria.

Strikingly, semapimod-treated Mtb exhibits significant upregulation of genes that encode for proteins involved in MCC such as methylcitrate synthase PrpC (19.48-fold), methylcitrate dehydratase PrpD (92.5-fold), the regulator of *prpC* and *prpD* genes– PrpR (28.55-fold), and an enzyme, Icl1 (68.69-fold). Icl1 of Mtb exhibits both the isocitrate lyase activity involved in the glyoxylate shunt, and methylisocitrate lyase activity which is responsible for catalysing the last reaction of the MCC, resulting in the production of succinate and pyruvate that are routed into the TCA cycle^46^. Although, MCC is responsible for metabolizing the toxic intermediates produced from catabolism of odd-chain fatty acids (OCFAs) such as cholesterol, which is the primary source of nutrient during host infection, it has been observed that both the glyoxylate shunt and MCC are functional in wild-type Mtb, even in the presence of glycolytic carbon source^47^. Icl1-deficient Mtb undergoes a progressive depletion of TCA cycle intermediates, which in-turn impair Mtb’s respiratory activity. Although, excess of leucine in the extracellular environment is lethal, intracellular leucine production is critical as it provides an alternative pool for acetyl-CoA biosynthesis which feeds into the respiratory and lipid metabolic pathways^48^. We believe that decline in intracellular leucine together with reduced expression of genes involved in its catabolism (*accD1*, *bkdA* and *bkdB*) triggers the induction of alternate pathway such as MCC to maintain the cellular acetyl-CoA pool.

The complex cell-wall lipids of Mtb provide a hydrophobic barrier around the bacterium. These can be esterified with multiple methyl-branched (MB) long chain fatty acids such as PDIM, sulfolipid-1, and di-, tri-, and poly-acyl trehalose. These MB lipids are synthesized by individual polyketide synthase (PKS) complexes requiring malonyl-CoA and methyl malonyl-CoA (MMCoA), which are derived from acetyl-CoA and propionyl-CoA, respectively. PDIM is an important virulence factor, and controls the movement of nutrients and large molecule inhibitors such as vancomycin across the bacterial cell membrane. While genes involved in synthesis of phthiocerol moiety of PDIM such as *ppsB*, *ppsC* and *ppsD* are downregulated, those involved in biosynthesis of SL-1 (*papA1* and *pks2*) and poly-acyl trehalose (*pks3*, *pks4*, *papA3*, and *fadD21*) are highly upregulated in drug-treated bacteria. Since, both PDIM and SL-1 biosynthetic pathways share a common precursor, MMCoA, it has been observed that reduction of PDIM synthesis favours SL-1 production due to increased flux of MMCoA to the SL-1 biosynthetic pathway, and *vice versa*^49^. Intrigued with these observations, we speculate that semapimod induces remodelling of cell-wall lipids, restricting L-leucine uptake by an unknown mechanism (Fig. 8). Targeting of PpsB by semapimod further supports the above hypothesis and warrants future studies to identify alterations in the cell envelope which impose restriction on L-leucine transport.

Uptake of host-derived nutrients including amino acids supports bacterial viability during infection^40, 41^. Targeting the amino acid transport system offers a novel strategy to curtail the supply of host-derived nutrients to the pathogens, which are metabolically quiescent during intracellular stay^50^. Semapimod, which has been found to inhibit the L-leucine uptake in Mtb, may not be the preferred choice of L-leucine inhibitor owing to its anti-inflammatory effect. It inhibits production of proinflammatory cytokine such as TNF-α, IL-1β, and IL-6. Also, it has been reported that semapimod interferes with the TLR4 signalling^17, 18, 19, 20^. These cytokines play a critical role in providing protection to the host against active TB infection^51^. Nonetheless, despite its anti-inflammatory activity which may promote bacterial proliferation, semapimod is able to reduce the intracellular burden of virulent Mtb H37Rv strain in lungs and spleen of the infected mice (Fig. 7). These findings provide an indirect evidence of the importance of L-leucine uptake for Mtb virulence during host infection.

Advancements in the genetic manipulation tools such as CRISPR-Cas technology^52^ has significantly contributed to enhancing our understanding of metabolic networks in mycobacteria. Screening of inhibitors against other Mtb auxotrophic strains is expected to provide novel insights into transport mechanisms of essential metabolites in Mtb that can be explored as novel anti-TB drug targets.

## Materials and Methods

### Strains and culturing of bacteria

For propagation of plasmids, *Escherichia coli* strain XL1Blue^TM^(Agilent) was used, whereas expression and purification of PpsB was performed in *E. coli* BL21 DE3 (Novagen). Mtb H37Rv was received from Dr. Ramandeep Singh at THSTI, India, and Mtb mc^2^ 6206 strain was provided by Dr. William Jacobs at Albert Einstein College of Medicine, NY, USA. *E. coli* was cultured in the Luria-Bertani medium (Becton Dickinson), Mtb H37Rv was cultured in Middlebrook 7H9 containing 0.05% tyloxapol (Merck), 0.5% glycerol (Merck) and 1X OADS (oleic acid-albumin-dextrose-saline) (7H9-O) or Middlebrook 7H11 containing 0.5% glycerol (Merck) and 1X OADS (7H11-O). For culturing Mtb mc^2^ 6206 strain, media were supplemented with 50µg/ml L-leucine (Merck) and 24µg/ml pantothenate (Merck) (7H9-PLO). Liquid cultures were grown either in Corning 50 mL centrifuge tubes (Corning) or in 250ml flasks (Corning) with not more than one-third volume of bacterial cultures, whereas plates were incubated at 37°C. Wherever required, we used 50µg/ml kanamycin (Merck), 50µg/ml ampicillin (Merck), and 150µg/ml hygromycin (Invitrogen) for *E. coli*, whereas for Mtb 25µg/ml kanamycin and 50µg/ml hygromycin were used.

### Screening of inhibitors

Repurposed library comprising of 3614 FDA-approved drugs was obtained from MedChemExpress (www.medchemexpress.com) in the 96-well plate format. All compounds were suspended in DMSO (Merck) at 5mM concentration, and stored frozen at -80C before use. For initial evaluation of compounds, 100µl Mtb mc^2^ 6206 culture, diluted to OD_600_ of ∼0.01 in 7H9-PLO medium was aliquoted in each well of the round-bottom 96-well plate containing 1µl of respective drugs, so that final conc. of drug became 50µM. Side wells were filled with 100µl medium to maintain humidity and to prevent loss of samples from drug-containing wells. As an untreated (UT) control, 100µl bacterial culture was treated with 1µl DMSO. Plates were incubated at 37 °C for 2-3 weeks until UT wells exhibited substantial growth. Drugs showing complete inhibition of growth by visual inspection of the pellet formation at the bottom were considered inhibitors of Mtb mc^2^ 6206, and used for determination of drug concentration inhibiting 90% growth (MIC_90_).

### Determination of MIC_90_

The MIC_90_ of active inhibitors was determined by using the resazurin assay. Briefly, Mtb mc^2^ 6206 was incubated at OD_600_ of 0.005 with different concentrations of drug ranging from 25-0.10µM in the flat bottom 96-well plates. After 2 weeks of incubation at 37 °C, cultures were transferred in the black 96-well plates and incubated with 10% of Alamar Blue^TM^ reagent (Thermo Scientific) for 16hrs 37 °C. Viability was calculated by comparing fluorescence of the drug-treated and DMSO-treated samples at 560nm of excitation and 590nm of emission. The minimum drug concentration at which ∼90% growth inhibition is found, was considered as MIC_90_.

### Macrophage infection and intracellular CFU analysis

For macrophage infection, THP1 cells were seeded in 24-well plates at a density of 2 × 10^5^ cells per well and differentiated using 20nM phorbol 12-myristate 13-acetate (PMA). After 48hr of differentiation, cells were washed once with incomplete RPMI medium (Himedia). Thereafter cells were infected with Mtb mc^2^ 6206 at 1:5 MOI (macrophage: bacteria) using RPMI medium containing 10% FBS medium, 24µg/ml pantothenate and 50µg/ml L-leucine (RPMI-PL). After 4hr of incubation, cells were washed with 1x phosphate buffered saline (1x PBS) for three times and replenished with RPMI-PL. After overnight incubation, cells were treated with 0.1, 0.2 and 1.0µM semapimod, and 2.5µM INH in four wells each and remaining 4 wells were left untreated. Drug treatment was repeated again after 3 days of incubation. On the 6^th^ day, cells were lysed in 0.1% Triton X100, and serial dilutions of the lysates were plated on 7H11-OADS-PL agar plates for CFU enumeration.

### Global gene expression analysis by RNA sequencing

Freshly inoculated culture of Mtb mc^2^ 6206 was treated with 50nM of semapimod at 0.40 OD, whereas another set was left untreated. After 24hrs of incubation at 37 °C with shaking at 200rpm, both semapimod-treated and untreated cultures were pelleted down, washed twice with 1x PBS and stored at - 80 °C before RNA extraction. Total RNAs were extracted from three biological replicates of untreated (control) and semapimod treated cultures followed by treatment with DNase I, as described previously^53^. The DNase-treated RNA samples were supplied to Clevergene (https://clevergene.in/) for further processing and sequencing, typically as reported earlier^53^. Genes that exhibit absolute log2 fold change ≥1, and FDR-adjusted *p* value of ≤0.01 were considered significant, and their expression profile is presented in the volcano plot and heatmap.

### Quantitative reverse transcription real-time PCR (qRT-PCR)

Total RNA was treated with DNase I (Ambion), and 500ng of DNase-treated RNA samples were subjected to complementary DNA (cDNA) synthesis using SuperScript IV reverse transcriptase, as recommended by the manufacturer (Thermo Scientific). qRT-PCR was performed by using 50ng cDNA and gene-specific primers (Supplementary Table 1) with the help of SYBR Green PCR master Mix (ABI). Amplification was monitored in real-time and quantification was carried out using ABI 7500 Fast Real-Time PCR System (Applied Biosystems), as mentioned earlier^53^.

### Measurement of ATP

Mtb mc2 6206 was cultured in the presence or the absence of 50nM semapimod for 24 h. For intracellular ATP estimation, 1ml of untreated and treated cultures at OD_600_ of 0.1 were pelleted by centrifugation, washed twice with 1x PBS and lysed in 200µl of 1x PBS by boiling at 98 °C for 10min. The bacterial lysates were centrifuged at 14,500 rpm for 3min to remove debris, and supernatant was collected for ATP estimation by using Bac Titer- Glo^TM^ assay kit as recommended by the manufacturer (Promega).

### Estimation of intracellular amino acids

Intracellular level of amino acids was estimated by liquid chromatography mass spectrometry (LC-MS/MS), as follows:

**a) Preparation of samples:** For estimation in the presence of semapimod, freshly-grown Mtb mc^2^ 6206 cultures were diluted to 1.0 OD_600_ and divided into two portions, one treated with 50ng/ml semapimod and the other one left untreated. After 24 hours of incubation at 37 °C with shaking at 200rpm, cultures equivalent to 5.0 OD_600_ were drawn from both the samples and pelleted by centrifugation. To measure the levels of amino acids under regular conditions in the WT or the Sem^R^ strains of Mtb mc^2^ 6206, bacteria were freshly inoculated as 0.1 OD_600_, and cultured for 7 days at 37 °C with shaking at 200rpm. Cultures equivalent to 5.0 OD_600_ were subsequently subjected to pelleting by centrifugation. After through washing with 1 x PBS, bacterial metabolites were extracted using methanol precipitation method, and subjected to LC-MS/MS.
**b) LC-MS/MS:** QTRAP® 6500+ MS system coupled with Ultra-high-performance liquid chromatographic system, (SCIEX ExionLC^TM^) was used for targeted LC-MS analysis. The ACQUITY UPLC HSS T3 column (1.8µm, 2.1 x 100mm; Waters) was used for separation, and the column oven temperature was set at 40 °C. Solvent A (10mM Ammonium formate with 0.1% formic acid in water) and solvent B (0.1% formic acid in methanol) were used for LC gradient with changing the concentration of solvent B as follows: 0 min, 1% B; 1 min, 15% B; 4 min, 35% B; 7 min, 95% B; 9 min, 95% B; 10 min, 1% B, 20 min, 1% B. The flow rate was kept at 300 µl/min and the injection volume was 5 μl. Following parameters were used for electrospray ionization: electrospray voltage, +5500 V; ion source temperature, 500 °C; curtain gas of 24, CAD gas 9, and gas 1 and 2 of 60 and 40 psi, respectively.
**c) Compound parameter:** Following optimized MRM parameters of de-clustering potential (DP), collision energies (CE), entrance potential (EP), and collision cell exit potential (CXP) were used: DP-40, EP-10, CE-15 & CXP-22. Commercially procured ultrapure L-leucine, valine and proline (Merck) were used for selection of Q1 and Q3 masses. Sharp peaks of pure compounds were chosen as references to identify the corresponding peaks in samples. Data analysis for LC-MS/MS was performed using the SCIEX (ANALYST 1.7.2) software. Analyte was confirmed by comparing the retention time and the ratio of characteristic transition between the test sample and the reference.

### Estimation of percent viability in response to vancomycin or semapimod treatment

Bacterial cultures were incubated at OD_600_ of 0.005 with different concentrations of vancomycin or semapimod in the flat bottom 96-well plates. After 2 weeks of incubation at 37 °C, growth was estimated by measuring OD_600_ of the cultures using multi-mode plate reader, which was used for measuring percent viability in the drug-treated cultures with respect to the untreated control.

### Cloning and expression of genes in Mtb

The *leuC-leuD* and *panC-panD* open reading frames (ORFs) were PCR amplified using Mtb H37Rv genomic DNA with gene-specific primer pairs (Supplementary Table 1) and GoTaq polymerase, as recommended by the manufacturer (Promega). PCR amplicons with *Nde* I and *Hind* III overhangs were gel purified and subjected to restriction digestion with the respective enzymes. The digested DNA fragments were column-purified and annealed at the same sites in a Hyg^R^ replicative *E. coli*-mycobacterium shuttle plasmid, pHsp60 under a constitutive promoter of *hsp60*^54^. The resulting recombinant plasmids, pHsp-*leuCD* and pHsp-*panCD* were electroporated in Mtb mc^2^ 6206 after sequence verification by Sanger’s sequencing. To ascertain expression of the respective genes, recombinant clones obtained by transformation with pHsp-*leuCD* were selected on 7H11-OADS agar plates containing pantothenate and hygromycin, whereas those obtained with pHsp-*panCD* were selected on 7H11-OADS agar plates containing L-leucine and hygromycin.

For expression of *ppsB*, the ∼4.6kb *ppsB* ORF was PCR amplified using Mtb H37Rv genomic DNA with Pr. 2396-2428 (PCR-A) and 2429-2397 (PCR-B) (Supplementary Table 1). Both the PCR-A and PCR-B fragments of ∼2.2 and ∼2.4 kb, respectively, were subsequently cloned in the pGEMT-easy plasmid by following the TA cloning method, as suggested by the manufacturer (Promega). Both the recombinant clones were sequence verified, which revealed the presence of PCR-A in the forward orientation in pGEMT-A clone and PCR-B in the reverse orientation in pGEMT-B clone. Both the clones were subjected to restriction digestion with *Nde I* and *Sma I*. PCR-A fragment of ∼2.2kb was gel eluted and cloned in the pGEMT-B plasmid at the same sites, to create full-length ppsB of ∼4.6kb. Next, DNA fragment comprising of full-length *ppsB* ORF was obtained from the positive clone of pGEMT-*ppsB* by *Nde I-Hind III* digestion, and cloned in pHsp60 at the same sites. To express *mce1D* and *ppe60*, the respective ORFs were PCR amplified from Mtb H37Rv genomic DNA using Pr. 2277-2278 and Pr. 2281-2282, respectively, (Supplementary Table 1), restriction digested and cloned in pHsp60 at *Nde I-Hind III* sites, as described above. The resulting recombinant plasmids– pHsp-*ppsB*, pHsp-*mce1D* and pHsp-*ppe60* were verified by Sanger’s sequencing and electroporated in wild-type or Sem^R^ Mtb mc^2^ 6206. The transformants were selected on 7H11-PLO agar plates containing hygromycin, and positive clones verified by colony PCR were used in subsequent experiments.

### Generation of Sem^R^ strain

To induce semapimod-resistance, 5ml culture of Mtb mc^2^ 6206 at OD_600_ of ∼2.0 was pelleted by centrifugation followed by suspension in ∼500µl 7H9-PLO medium. Aliquots of 50µl of the culture suspension were spread on multiple 7H11-PLO agar plates containing 100nM semapimod. After 4-8 weeks of incubation at 37 °C, single colonies appeared on drug-containing plates were cultured in 1ml 7H9-PLO medium. Subsequently, cultures of the wild-type and putative Sem^R^ strain of Mtb mc^2^ 6206 were spotted on 7H11-PLO agar plates with or without 50nM semapimod to monitor growth.

### Genomic DNA extraction from mycobacteria and whole genome sequencing

Mtb mc^2^6206 wild type and the Sem^R^ strains were grown in 7H9-PLO, and genomic DNA extraction was performed using the conventional cetyltrimethylammonium bromide (CTAB)-based method. Whole genome sequencing (WGS) of the Mtb strains was conducted using the PacBio Sequel II long read sequencing platform. The WGS was outsourced to Nucleome Informatics Private Limited, Hyderabad, India. A detailed sequencing workflow was employed to ensure the generation of high-quality data. Briefly, DNA quantification was performed using Qubit fluorometry, while DNA quality was assessed on a 1% agarose gel. Purity ratios were analyzed using the NanoDrop 2000 spectrophotometer. DNA fragments between 7–10 kb were prepared for SMRTbell library creation utilizing the Megaruptor 3 system, followed by purification with AMPure PB beads. The Agilent FEMTO Pulse analyzer was used to verify size distribution, and DNA concentration was measured with the DNA HS assay on the Qubit 3.0 fluorometer. Libraries were constructed with the Template Prep Kit, starting with the removal of single-strand overhangs, followed by DNA damage repair and End-Repair/A-tailing. Adapters were then ligated to the repaired DNA fragments to form SMRTbell libraries, with hairpin dimers removed using AMPure PB beads. The final pooled libraries were further purified and analyzed with the Agilent FEMTO Pulse. Primer annealing and polymerase binding were conducted using the Sequel II binding kit 2.2 before loading the libraries onto SMRTcells for sequencing in CCS/HiFi mode on the PacBio Sequel II system. This robust workflow ensured the production of high-quality sequencing data essential for precise genomic analysis.

### Bioinformatic analysis for genome assembly preparation and polymorphism detection

PacBio Sequel II raw subreads were processed into high-fidelity (HiFi) reads using the ‘CCS’ tool (version 6.2.0). The genome assembly was constructed using ‘Trycycler’ (version 0.5.4) with its default parameters^55^. Three individual assemblies were initially created with the tools ‘Flye’ (version 2.9.2), ‘Minipolish’ (version 0.1.3), and ‘wtdbg2’ (version 2.5)^56, 57, 58^. These assemblies were combined to generate a consensus assembly using Trycycler’s default configuration. Genome completeness was evaluated using BUSCO (version 5.4.5), with the actinobacteria_class_odb10 lineage dataset as the reference^59^. For comparative genomics, variant calling was performed using Genome Analysis Toolkit (GATK, version 4.5), FreeBayes (version 1.3.7), and TB-Profiler (version 5.0.1)^60, 61, 62^. Mutations of interest were carefully selected based on their confidence levels to ensure accuracy.

### Cloning, expression and purification of PpsB

Full-length *ppsB* ORF fragment was obtained from pGEMT-*ppsB* by *Nde I-Hind III* digestion and cloned at the same sites in pET28b (Novagen) to express the recombinant protein with 6x His tags at the N-terminus *in E. coli*. For this, pET28b-*ppsB* was transformed in *E. coli* C43(DE3) cells and recombinant clones were selected on LB agar containing kanamycin. Single colony of *E. coli*::pET28b-*ppsB* was inoculated in 10ml of LB broth medium containing kanamycin and culture was grown to turbidity at 37 °C with shaking at 200rpm. Secondary culture was subsequently inoculated from the seed culture in one liter of LB broth medium with kanamycin. Expression of 6XHis-PpsB was induced by adding 1mM IPTG in the culture at OD_600_ of 0.80. After overnight incubation with IPTG at 18 °C, culture was pelleted and washed twice with 1x PBS. Induced culture pellet was suspended in 50ml of lysis buffer (50mM Tris, pH 8.0, 50mM NaCl and 5% glycerol) and cell lysis was performed by using French press. Cell debris was removed by centrifugation and clarified cell lysate was incubated with Ni-NTA beads (Qiagen) for 3 h at 4 °C. Beads were subsequently transferred into an empty column and unbound proteins were allowed to pass through. Beads immobilized with 6XHis-PpsB were washed with 5 bed volumes of the lysis buffer, followed by washing with 5 bed volumes of wash buffer (lysis buffer containing 20mM imidazole). Immobilized 6XHis-PpsB was subsequently eluted with elution buffers (lysis buffer containing 100-300mM imidazole). The purity of eluted protein in different elution fractions was assessed by SDS-PAGE. Fractions showing >90% purity of PpsB were pooled and dialyzed against lysis buffer. The purified proteins were stored in small aliquots at -80°C for subsequent use.

### Protein-drug interaction by BLI-Octet

Optical interference-based Bio layer interferometry (BLI) from Octet system by ForteBIO was used to ascertain the interaction of semapimod with Mtb PpsB. Briefly, purified 6xHis-tagged PpsB protein was dialyzed in 10 mM sodium acetate pH 3.9, followed by immobilization onto the amine reactive group sensor (2nd-generation) (AR2G) up to a level of 2.1 nm. Different concentrations of semapimod were prepared in water and used to acquire differential graded response. Binding constant was calculated as per the standard steps of baseline (60 s), association (180 s) and dissociation (180 s).

### Extraction of lipids

To obtain the apolar lipids, bacterial cultures equivalent to OD_600_ of ∼150 were pelleted, washed with 10ml of water and suspended in 2 mL methanol-0.3% NaCl (10:1) and 1 mL petroleum ether. After mixing vigorously to emulsify the organic and the aqueous layers, samples were centrifuged at a speed of 2000 x *g* for 5 minutes to allow mixture to separate. The upper organic layer was collected in the microcentrifuge tubes, and the extraction was repeated by adding 1 mL petroleum ether to the lower aqueous phase and stirred for another 30 min. The organic layer was collected as described above and combined with the previous extracts. Petroleum ether was subsequently evaporated by vacuum drying at the room temperature and the resulting apolar lipids were stored at -80 °C for analysis by thin layer chromatography (TLC).

To prepare MAMES, culture pellets equivalent to OD_600_ of ∼150 were suspended in 1ml of 20% tetrabutylammonium hydroxide and incubated at 100 °C for overnight. After overnight incubation, 0.4 mL of dichloromethane and 50µl of iodomethane were added to the reaction mixture and stirred for 1 h at room temperature. The biphasic mixture was separated by centrifugation at 2000 x *g* for 5 minutes, and lower organic layer containing MAMES was collected in the microcentrifuge tubes. The MAMES fraction was completely dried and suspended in 1 mL of diethyl ether. The supernatant was collected in a new tube and vacuum dried. The dried fraction was suspended in 0.2ml of toluene and 0.1ml of acetonitrile, and after complete dissolution 0.2ml more acetonitrile was added. Samples were chilled at -20 °C for overnight and the MAMES were collected by centrifugation at 10000 x *g* for 20 minutes. MAMES present in the pellet fraction were stored at -80 °C for further analysis by TLC.

### Analysis of lipids by TLC

Both the apolar lipids and MAMES were detected on aluminum-backed silica gel 60F254 plates (Merck). Dried lipid pellets were suspended in dichloromethane and equal volume of samples from both the strains were spotted on the TLC plates. Apolar lipids were resolved by two dimensional-TLC (2D-TLC) using petroleum ether:ethyl acetate solvent (98: 2) for 3 times in the first dimension, and petroleum ether:acetone mixture (98: 2) for one time in the second dimension. MAMES were resolved by one dimensional-TLC using petroleum ether:diethyl ether (95:5) solvent for 6 times. After complete run, TLC plates were air dried and sprayed with 5% phosphomolybdic acid solution, prepared in ethanol. Lipids were visualized on the TLC plates after charring at 100 °C for 2-5 minutes.

### Mice infection

Mice infection was performed as per the guidelines by the committee for the purpose of control and supervision of experiments on animals (CPCSEA, India). Animal infection experiment was performed after receiving approval from the Institutional Animal Ethics Committee of THSTI. Briefly, 6- to 8-week-old female BALB/c mice were infected with single cell suspension of mouse-passaged Mtb H_37_Rv at ∼5×10^7^ cells/ml via aerosol route in a closed aerosol chamber in the biosafety level 3 laboratory. To check the establishment of infection, bacterial load in lungs were examined at day 1 post-infection and the disease progression was assessed at day 21 post-infection, by CFU plating on selective 7H11-OADS agar plates containing 100µg/ml carbenicillin, 22µg/ml trimethoprim, 22µg/ml amphotericin B, 11µg/ml vancomycin, 27.5µg/ml polymyxin B and 0.88µg/ml cycloheximide. At day 21, mice were divided into two groups; one group was given semapimod (5mg/kg in 5% sucrose) by daily oral gavaging whereas the other half was given equal dose of 5% sucrose. CFUs were enumerated at day 28 post-semapimod or sucrose treatment to examine bacterial burden in lungs, and in spleen for dissemination.

### Statistics and reproducibility

Some experiments were only performed with 2 biological replicates. Statistical analysis was performed with data obtained from 3 or more biological repeats by determining *p* values with the help of GraphPad Prism version 10.3.1 (464) software.

## Data availability

Raw data of the RNASeq are available from the GEO database under project accession number GSE284673. Raw data of the whole genome sequence of Mtb mc^2^ 6206 can be accessed from the Sequence Read Archive (SRA) database under the Bio-project PRJNA1118666, which can be reviewed by using the following link:https://dataview.ncbi.nlm.nih.gov/object/PRJNA1118666?reviewer=i6d7semeju6p35ks3vnp6gvda3.

## Acknowledgements

We thank Dr. Ramandeep Singh at the Translational Health Science and Technology Institute, India, Dr. Rajesh Gokhale at the National Institute of Immunology, India and Dr. William Jacobs at the Albert Einstein College of Medicine, USA for providing us the mycobacterial strains. Ms. Shivani Goswami (Lab Technician) and Mr. Manish Bansal (Technical Officer) are acknowledged for providing assistance in the screening of inhibitors. We acknowledge technical support by the research staffs at the Infectious Disease Research Facility (IDRF) at THSTI in the animal studies. Experimental Animal Facility (EAF) at THSTI is acknowledged for providing animals. This work was supported by the core research funding from THSTI. Research fellowships to L.A. (Award No. 15/12/2019 (ii) EU-V) and MYK (Award No. 15/12/2019 (ii) EU-V) by Council of Scientific & Industrial Research (CSIR), and to H.G from Science and Engineering Research Board, Department of Science and Technology, Govt. of India (Award No. RJF/2022/000040) are acknowledged.

## Author information

### Author contributions

N.A. designed the research, performed the experiments, analysed the data, wrote the paper and provided overall supervision of the study. H.G., Eeba, L.A., and M.Y.K. performed the experiments and provided inputs in manuscript writing. S.K.B. and B.D. analysed genome sequence data.

