## Supplementary Information for "Identifying a novel mechanism of L-leucine uptake in *Mycobacterium tuberculosis* using a chemical genomic approach"

**Supplementary Figure 1. Characteristics of the small molecules' library. a**, Bar graph shows status of molecules in the library acting on different metabolic pathways. **b**, Pi chart depicts various diseases that are targeted by inhibitors. **c**, Shown is the ranking of FDA-approved molecules as per the different clinical trial phases.

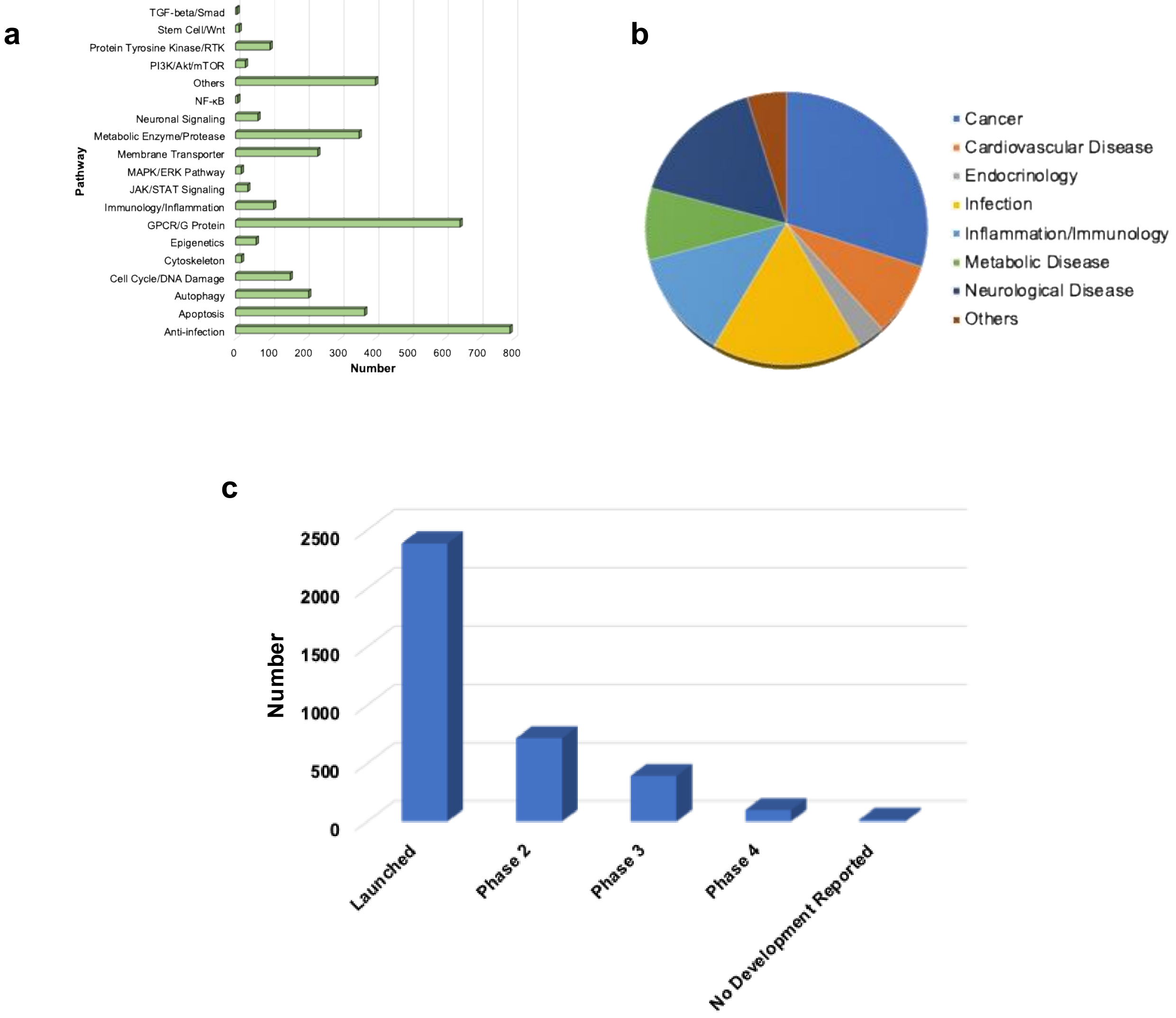

**Supplementary Figure 2. Validation of RNASeq data by qRT-PCR.** Bar graph shows the expression levels of representative genes in semapimod-treated bacteria, estimated by qRT-PCR. Fold-change was calculated with respect to untreated samples after normalization with *rrs*, which remained consistent across the two groups. Mean±s.d. values of n=2 biological replicates are shown.

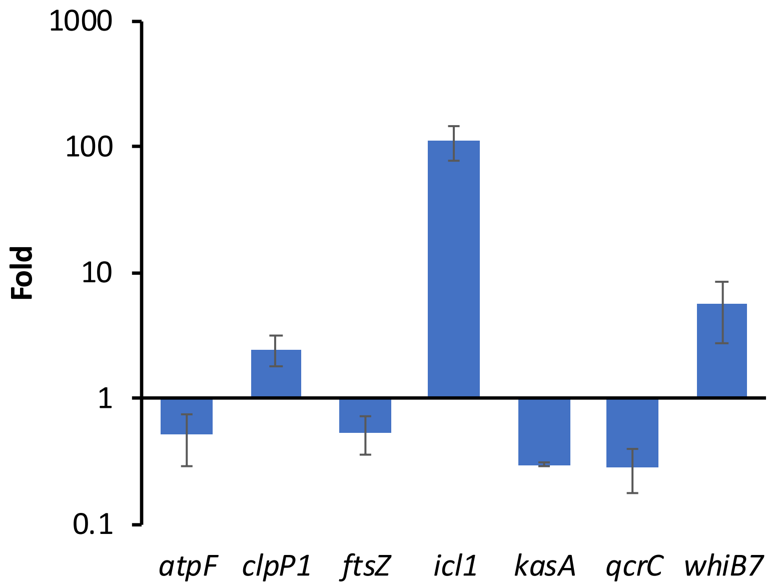

**Supplementary Figure 3: Semapimod treatment causes reduction in intracellular ATP level.** Intracellular ATP was estimated in UT and treated Mtb mc² 6206 exposed to 50nM semapimod (Sem), by using Bac TiterGlo™ assay kit, as suggested by the manufacturer (Promega). Bar graph shows % change in the intracellular ATP levels following drug exposure. Mean±s.d. values of n=2 biological replicates are shown.

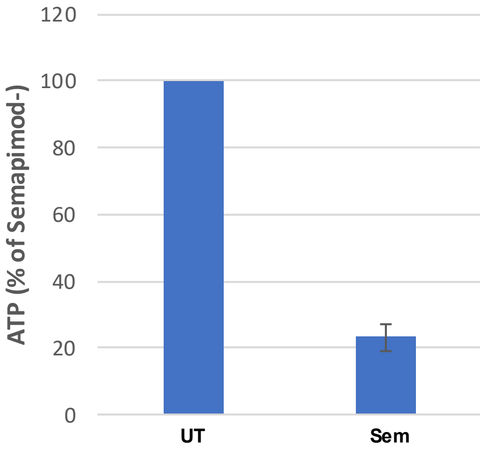

**Supplementary Figure 4. Effect of withdrawal of L-leucine on expression of semapimod-regulated genes in Mtb mc[2] 6206.** Bar graph shows the expression level of various genes in the presence or the absence of metabolites, as estimated by qRT-PCR. PL+, 7H9-PLO; PL-, 7H9-O; P-, 7H9-LO; L-, 7H9-PO. Fold-change in expression was calculated in all the samples with respect to PL+ after normalization with *sigA*, which remained consistent in all the samples. Mean±s.d. values of n=2 biological replicates are shown. P values were calculated using Graphpad prism version 10.

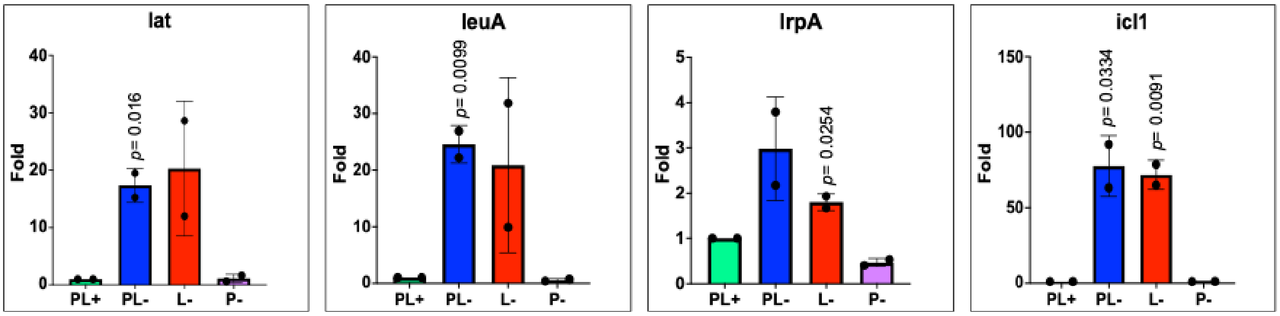

**Supplementary Figure 5. Effect of Semapimod treatment on the intracellular levels of valine and L-proline in the Mtb mc$^2$ 6206.** Intracellular valine and L-proline were estimated in untreated (Control) and semapimod-treated (Treated) bacteria, after 24 hours of drug treatment, by mass spectrometry as described in Materials and Methods. Mean+s.d values from n=2 replicates are shown.

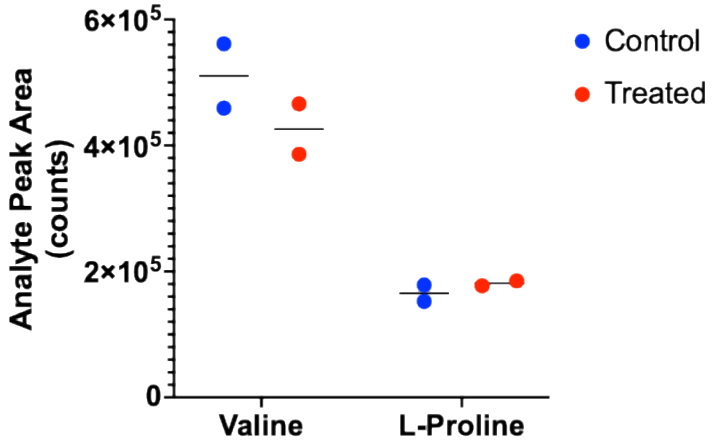

**Supplementary Figure 6. List of *M. smegmatis* genes potentially involved in L-leucine transport.**

| Gene | Description |
|---|---|
| MSMEG_1216 | ABC-type transport system periplasmic substrate-binding protein |
| MSMEG_1217 | ABC-type transport system ATP-binding protein I |
| MSMEG_1218 | ABC-type transport system ATP-binding protein II |
| MSMEG_1219 | ABC-type transport system permease protein II |
| MSMEG_1220 | ABC-type transport system permease protein I |
| MSMEG_3247 | Branched-chain amino acid ABC transporter substrate-binding protein |
| MSMEG_3248 | ABC transporter branched chain amino acid transport permease |
| MSMEG_3249 | Branched-chain amino acid ABC transporter, permease protein |
| MSMEG_3250 | ABC transporter, ATP-binding protein |
| MSMEG_3251 | Branched-chain amino acid ABC transporter ATP-binding protein |
| MSMEG_6876 | Branched chain amino acid transport ATP-binding protein |
| MSMEG_6877 | Branched-chain amino acid transport system ATP-binding protein |
| MSMEG_6878 | Inner-membrane translocator |
| MSMEG_6879 | Integral membrane protein of the ABC-type Nat permease for neutral amino acids NatD |
| MSMEG_6880 | Hydrophobic amino acid ABC transporter, putative |

**Supplementary Figure 7. Estimation of intracellular L-leucine and valine in the Mtb mc$^2$ 6206.** Intracellular L-leucine and valine were estimated in the wild-type (WT) and Sem$^R$ bacteria, after 7 days of culturing, by mass spectrometry as described in Materials and Methods. Mean+s.d values from n=3 replicates are shown. P values were calculated using Graphpad prism version 10.

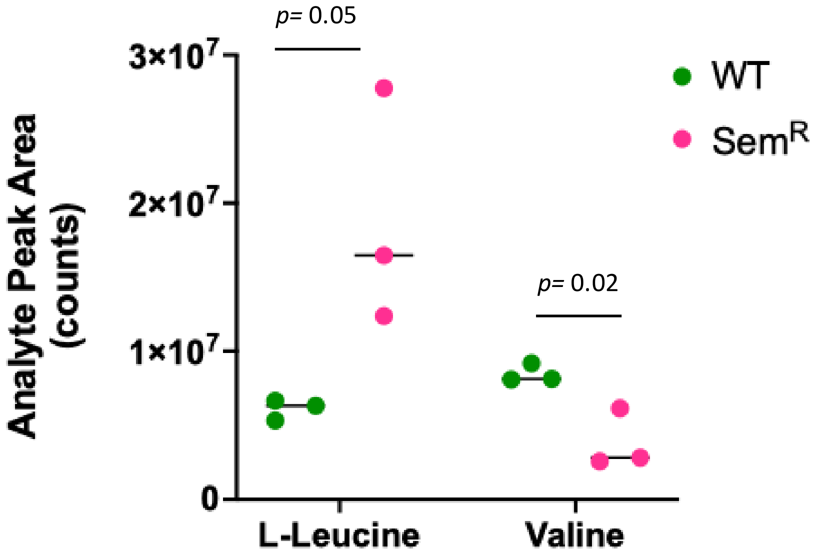

**Supplementary Figure 8. Analysis of $MIC_{90}$ of Rif and Inh against Semapimod$^R$ strain of Mtb mc$^2$ 6206.** $MIC_{90}$ of standard TB drugs, rifampicin and INH was determined against WT and Sem$^R$ strains using 96-well plate-based assay, as described in the Materials and Methods. As can be seen, resistance to semapimod does not alter bacterial susceptibility to either of the frontline TB drugs.

| | Rifampicin | INH |
|---|---|---|
| WT | 0.11µM | 0.40µM |
| Sem$^R$ | 0.11µM | 0.40µM |

**Supplementary Figure 9. Comparative sequence analysis of Ppe60 in the wild-type and Sem^R strains of Mtb 6206.** Sequence alignment of Ppe60 from WT and Sem^R strains was performed by Clustal Omega (https://www.ebi.ac.uk/jdispatcher/msa/clustalo) using default parameters. ‘*’ represents identical residues; ‘.’ represents weakly similar residues; ‘:’ signifies conserved residues with strongly similar properties, whereas completely dissimilar changes are shown by gaps.

```
SemR  VVDFGALPPEINSARMYAGPGSASLVAAAKMWDSVASDLFSAASAFQSVVWGLTTGSWIG 60
WT    VVDFGALPPEINSARMYAGPGSASLVAAAKMWDSVASDLFSAASAFQSVVWGLTVGSWIG 60
      ******************************************************.*****

SemR  SSAGLMVAAASPYVAWMSVTAGQAELTAAQVRVAAAAYETAYGLTVPPPVIAENRAELMI 120
WT    SSAGLMAAAASPYVAWMSVTAGQAQLTAAQVRVAAAAYETAYRLTVPPPVIAENRTELMT 120
      ******.*****************:***************** ************:***

SemR  LIATNLLGQNTPAIAVNEAEYGEMWAQDAAAMFGYAATAATATEALLPFEDAPLITNPGG 180
WT    LTATNLLGQNTPAIEANQAAYSQMWGQDAEAMYGYAATAATATEALLPFEDAPLITNPGG 180
      * ************ .*:* *.:**.*** **:***************************

SemR  LLEQAVAVEEAIDTAAANQLMNNVPQALQQLAQPTKSIWPFDQLSELWKAISPHLSPLSN 240
WT    LLEQAVAVEEAIDTAAANQLMNNVPQALQQLAQPAQGVVPSSKLGGLWTAVSPHLSPLSN 240
      **********************************::.: * .:*. **.*:*********

SemR  IVSMLNNHVSMTNSGVSMASTLHSMLKGFAPAAAQAVETAAQNGVQAMSSLGSQLGSSLG 300
WT    VSSIANNHMSMMGTGVSMTNTLHSMLKGLAPAAAQAVETAAENGVWAMSSLGSQLGSSLG 300
      : *: ***:** .:****:.********:************:*** **************

SemR  SSGLGAGVAANLGRAASVGSLSVPQAWAAANQAVTPAARALPLTSLTSAAQTAPGHMLGG 360
WT    SSGLGAGVAANLGRAASVGSLSVPPAWAAANQAVTPAARALPLTSLTSAAQTAPGHMLGG 360
      *********************** ************************************

SemR  LPLGHSVNAGSGINNALRVPARAYAIPRTPAAG 393
WT    LPLGHSVNAGSGINNALRVPARAYAIPRTPAAG 393
      *********************************
```

**Supplementary Figure 10. Overexpression of *mce1D* does not affect susceptibility of Mtb mc$^2$ 6206 to semapimod.** Effect of *mce1D* overexpression in the WT and Sem$^R$ strains on bacterial viability in the presence of semapimod. Viability of WT and Sem$^R$ strains constitutively expressing *mce1D* was assessed by 96-well plate-based assay, as described in Materials and Methods. Percent viability was calculated with respect to UT cultures after 2 weeks of exposure to different concentrations of inhibitors. Mean±s.d. values of n=2 biological and n=2 technical replicates are shown.

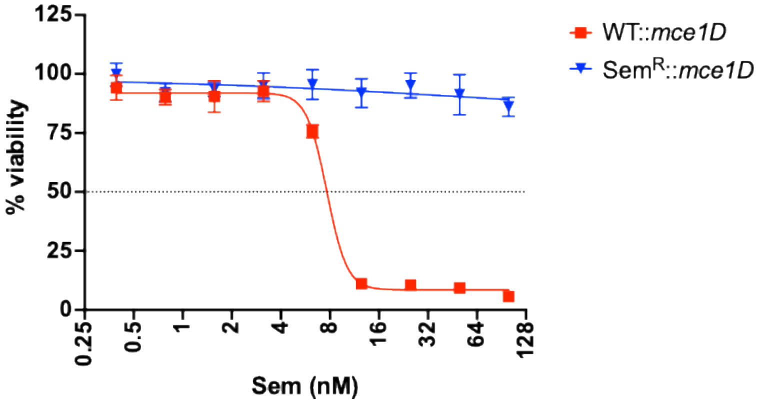

**Supplementary Figure 11. Overexpression of *ppe60* does not affect susceptibility of Mtb mc$^2$ 6206 to semapimod. a-b**, Effect of *ppe60* overexpression in the WT and Sem$^R$ strains on bacterial viability was determined in the presence of semapimod by 96-well plate-based assay, as described in Materials and Methods. Viability was assessed by visual inspection of the plate (**a**) as well as by determining percent viability (**b**). Percent viability was calculated with respect to UT cultures after 2 weeks of exposure to different concentrations of semapimod. Mean±s.d. values of n=2 biological replicates are shown.

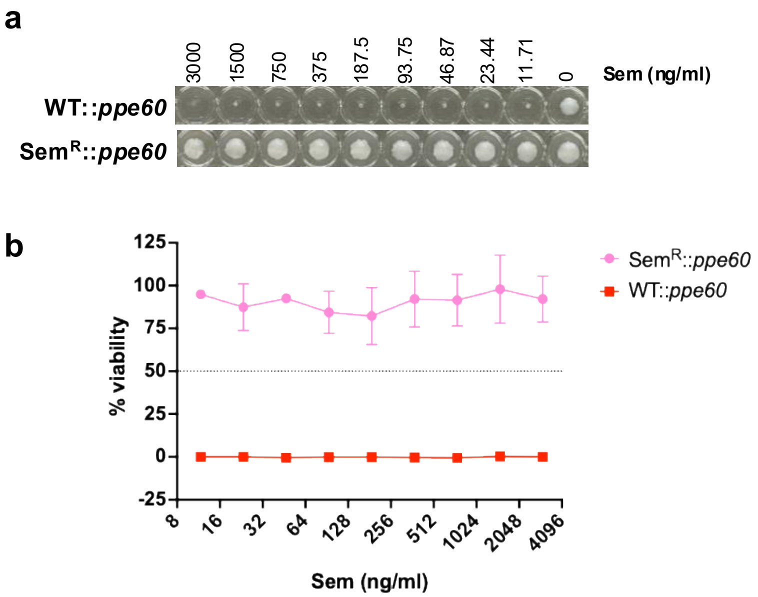

**Supplementary Figure 12. SDS-PAGE analysis of purified PpsB of Mtb.** Shown is the Coomassie Brilliant Blue-stained denaturing polyacrylamide gel with 10µg of purified PpsB with 6x His tags at the N-terminus. Molecular mass of the purified protein was determined by using the prestained molecular mass marker.

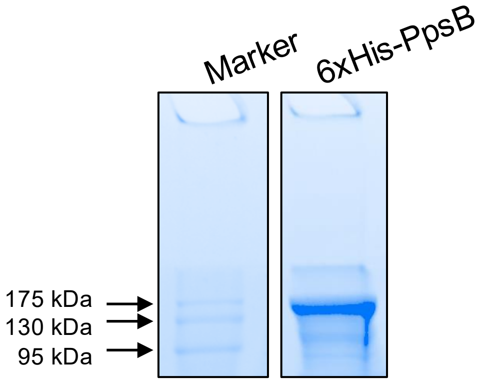

**Supplementary Figure 13. Comparative analysis of Cell wall mycolic acid profile of WT and Sem$^R$ strains of Mtb mc$^2$ 6206.** Cell wall mycolic acid methyl esters (MAMES) were extracted from both the strains and analyzed by one-dimensional TLC, as described in Materials and Methods. As can be seen, no change is observed in the levels of different MAMES between the two strains.

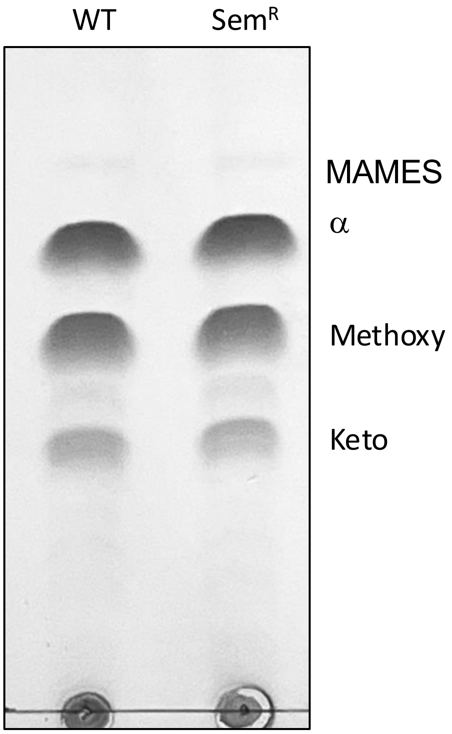

**Supplementary Figure 14. Susceptibility of Mtb H37Rv isolated from mice lungs post-semapimod treatment to vancomycin.** Susceptibility of Mtb H37Rv obtained from lungs of untreated (UT) and semapimod-treated (Sem) infected mice to vancomycin was assessed by 96-well plate-based assay, as described in Materials and Methods. Percent viability was calculated with respect to UT cultures after 2 weeks of exposure to different concentrations of vancomycin. Mean±s.d. values of n=2 biological replicates are shown.

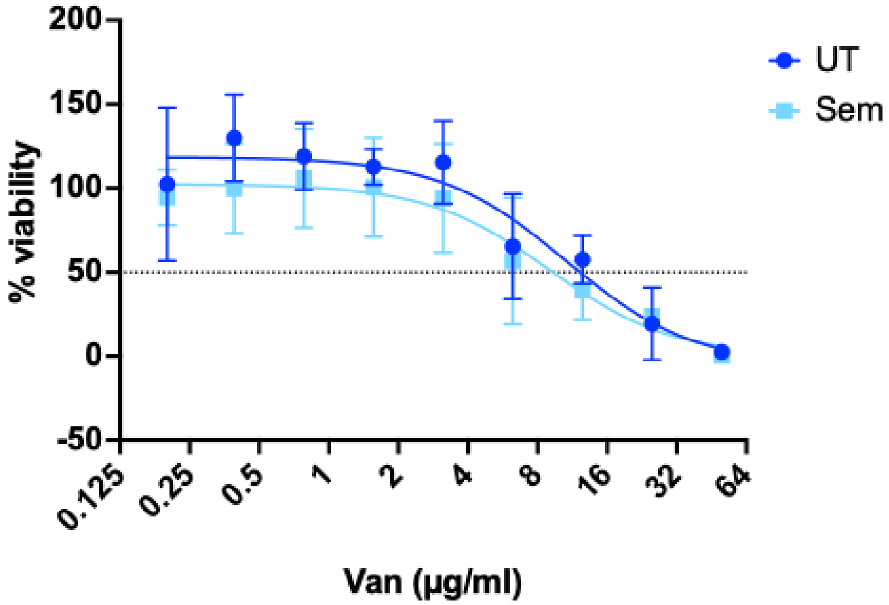

**Supplementary Table 1.** Shown is the list of oligonucleotides, used in this study.

| Oligo ID | Sequence (5'–3') | Description |
|---|---|---|
| atpF_RTF | GGTGAAGTGAGCGCGATTGTCC | Forward oligo used in qRT-PCR-based expression analysis of Mtb atpF |
| atpF_RTR | ACCATAGCGTCACGTTCCCG | Reverse oligo used in qRT-PCR-based expression analysis of Mtb atpF |
| clpP1_RTF | AAGTGACTGACATGCGTTCG | Forward oligo used in qRT-PCR-based expression analysis of Mtb clpP1 |
| clpP1_RTR | AGAGGCTGATGTCCTTGCTG | Reverse oligo used in qRT-PCR-based expression analysis of Mtb clpP1 |
| FtsZ_RTF | CCGGGTCTAATCAACGTCGACTTCG | Forward oligo used in qRT-PCR-based expression analysis of Mtb ftsZ |
| FtsZ_RTR | TCGACATCAGCACGCCTTGC | Reverse oligo used in qRT-PCR-based expression analysis of Mtb ftsZ |
| icl_RTF | GCTTCTACCGCACCAAGA | Forward oligo used in qRT-PCR-based expression analysis of Mtb icl1 |
| icl_RTR | TCGAGGTGCTTTTTCCAGT | Reverse oligo used in qRT-PCR-based expression analysis of Mtb icl1 |
| kasA_RTF | AAGTGGGATCTAGCGGTCAAGA | Forward oligo used in qRT-PCR-based expression analysis of Mtb kasA |
| kasA_RTR | ATCCTCTCGGCTCCACCTAGAC | Reverse oligo used in qRT-PCR-based expression analysis of Mtb kasA |
| qcrC_RTF | GTGTTGCTGCTGATAGCGCTG | Forward oligo used in qRT-PCR-based expression analysis of Mtb qcrC |
| qcrC_RTR | GATCAGACTCGGCCCGTGGTC | Reverse oligo used in qRT-PCR-based expression analysis of Mtb qcrC |
| whiB7_RTF | TGTCGGTACTGACAGTCCCC | Forward oligo used in qRT-PCR-based expression analysis of Mtb whiB7 |
| whiB7_RTR | TATCTCACCACCCCAAACGC | Reverse oligo used in qRT-PCR-based expression analysis of Mtb whiB7 |
| ppsB_RTF | CGGTTGCGGTGGTTGGCATC | Forward oligo used in qRT-PCR-based expression analysis of Mtb ppsB |
| ppsB_RTR | CCCGCGGTGTGATACCGAAG | Reverse oligo used in qRT-PCR-based expression analysis of Mtb ppsB |
| mce1D_RTF | CTGACGAACAACACGGTGGTCG | Forward oligo used in qRT-PCR-based expression analysis of Mtb mce1D |
| mce1D_RTR | CAACTGAATGTTCCGCGACGC | Reverse oligo used in qRT-PCR-based expression analysis of Mtb mce1D |
| rrs_RTF | TCCGGCCACACTGGGACTGAGATAC | Forward oligo used in qRT-PCR-based expression analysis of Mtb rrs |
| rrs_RTR | TATTACCGCGGCTGCTGGCAC | Reverse oligo used in qRT-PCR-based expression analysis of Mtb rrs |
| sigA_RTF | CCATCCCGAAAAGGAAGACC | Forward oligo used in qRT-PCR-based expression analysis of Mtb sigA |
| sigA_RTR | TCGAGGTCTGGTTCAGCGTC | Reverse oligo used in qRT-PCR-based expression analysis of Mtb sigA |
| Pr. 2396 | NNCATATGATGCGAACGGCTTTCAGCCGG | For PCR amplification of ppsB ORF |
| Pr. 2428 | ACCCGGGCGATCAACTCATCG | |
| Pr. 2429 | GCCGACCGAACAAATCGATG | |
| Pr. 2397 | NNAAGCTTCATTGTGTTCCTCTTAGTCGTTC | |
| leuC_ORF_F | NNCATATGGCCTTGCAGACCGGCGAG | For PCR amplification of leuC-leuD ORFs |
| leuD_ORF_R | NNAAGCTTTCAGGGGGCGGGTAGAGTGC | |
| panC_ORF_F | NNCATATGACGATTCCTGCGTTCCATC | For PCR amplification of panC-panD ORFs |
| panD_ORF_R | NNAAGCTTCTATCCCACACCGAGCCGGGGGTC | |
| Pr. 2277 | NNCATATGAGCACCATCTTTGATATCCGCA | For PCR amplification of mce1D ORF |
| Pr. 2278 | NNAAGCTTCATTGACCCCCTCCTGCCTCA | |
| Pr. 2281 | NNCATATGGTGGATTTCGGGGCGTTACC | For PCR amplification of ppe60 ORF |
| Pr. 2282 | NNAAGCTTCTATCCGGCGGCCGGTGTGC | |
